# Binding Position Dependent Modulation of Smoothened Activity by Cyclopamine

**DOI:** 10.1101/2024.02.08.579369

**Authors:** Kihong Kim, Prateek D. Bansal, Diwakar Shukla

## Abstract

Cyclopamine is a natural alkaloid that is known to act as an agonist when it binds to the Cysteine Rich Domain (CRD) of the Smoothened receptor and as an antagonist when it binds to the Transmembrane Domain (TMD). To study the effect of cyclopamine binding to each binding site experimentally, mutations in the other site are required. Hence, simulations are critical for understanding the WT activity due to binding at different sites. Additionally, there is a possibility that cyclopamine could bind to both sites simultaneously especially at high concentration, the implications of which remain unknown. We performed three independent sets of simulations to observe the receptor activation with cyclopamine bound to each site independently (CRD, TMD) and bound to both sites simultaneously. Using multi-milliseconds long aggregate MD simulations combined with Markov state models and machine learning, we explored the dynamic behavior of cyclopamine’s interactions with different domains of WT SMO. A higher population of the active state at equilibrium, a lower activation free energy barrier of *∼* 2 kcal/mol, and expansion of the hydrophobic tunnel to facilitate cholesterol transport agrees with the cyclopamine’s agonistic behavior when bound to the CRD of SMO. A higher population of the inactive state at equilibrium, a higher free energy barrier of *∼* 4 kcal/mol and restricted the hydrophobic tunnel to impede cholesterol transport showed cyclopamine’s antagonistic behavior when bound to TMD. With cyclopamine bound to both sites, there was a slightly larger inactive population at equilibrium and an increased free energy barrier (*∼* 3.5 kcal/mol). The tunnel was slightly larger than when solely bound to TMD, and showed a balance between agonism and antagonism with respect to residue movements exhibiting an overall weak antagonistic effect.

## Introduction

G-Protein Coupled Receptors (GPCRs) represent the largest family of human cell surface receptors that transmits signals across the cellular membrane. Upon binding of a ligand or through thermal fluctuations, the receptor undergoes a conformational change from an inactive state to an active state. This leads to heterotrimeric G-Protein binding at the intracellular membrane to initiate downstream signaling pathways. ^1^ Due to their crucial role in cellular signaling, GPCRs have become prime targets for drug development. In fact, 34% of all US Food and Drug Administration (FDA)-approved drugs target Class A and B GPCRs.^2^

Smoothened (SMO) is a member of the Frizzled (Fz) (Class F) family of GPCRs. It is composed of a heptahelical transmembrane domain (TMD), an extracellular cysteine-rich domain (CRD), and a linker domain (LD) that connects CRD and TMD. Smoothened plays a vital role in maintaining the Hedgehog (Hh) signaling pathway. Activation of Hh pathway begins with the binding of Hh ligands to the Patched (PTCH) receptor, causing the inhibition of PTCH and subsequent activation of SMO.^3,4^ The Hh pathway is crucial in ensuring stability during processes such as cell differentiation, regenerative responses in adults, and embryonic development.^5–8^ The dysregulation of this Hh pathway can lead to a wide range of diseases. Insufficient Hh activity is linked to birth defects such as holoprosencephaly and brachydactyly, while hyperactive Hh activity is associated with cancers such as basal cell carcinoma and medulloblastoma.^9,10^

Cyclopamine is a naturally occurring alkaloid found in corn lily (Veratrum californicum) that gained attention due to its association with birth defects in lambs.^11^ It hinders the separation of the embryonic brain into two lobes, resulting in a rare condition known as cyclopia, or a single eye disease.^12^ This unique effect of cyclopamine on fetal development led to its name, derived from the cycloptic lambs that were observed by Idaho lamb farmers in 1950s.^13^ It was only until the 2000s that cyclopamine’s behavior was explained by scientists, linking it to Hh inhibition.^14^ Cyclopamine gained significant attention due to its unique ability to inhibit the Hh signaling pathway by targeting the Smoothened (SMO) receptor.^14–16^ Its discovery has provided valuable insights into the mechanisms of Hh pathway regulation and has paved the way for the development of novel therapeutic strategies for Hh-dependent diseases, particularly cancer.^17^ Cyclopamine, through its inhibition of the Hh signaling pathway, shows promise as a treatment for various cancers with dysregulated Hh pathway activity.^17^ However, cyclopamine itself has toxic nature, and safer derivatives of cyclopamine are needed. ^17^ While Vismodegib, a synthetic derivative of cyclopamine, has shown remarkable success with FDA approval in January 2012 for treating metastatic or locally advanced basal cell carcinoma in adults,^18–20^ it is susceptible to chemoresistance,^21^ highlighting the need for continued research in this area.

Previous experiments have shown that cyclopamine can act as an antagonist by binding to and inhibiting at TMD site in SMO lacking CRD (SMOΔCRD).^14^ On the other hand, recent experimental results have shown that cyclopamine can increase Hh pathway activity by binding to CRD of mSmoD477G/E522K.^22^ The agonistic activity of cyclopamine in this case, however, does not reach the same level as Shh-based activation, making it a partial agonist. Determining the true effect of cyclopamine on wild-type hSMO is challenging, as experimentally probing the activity of one site requires mutation or deletion on the other site. Additionally, especially at saturated concentrations of cyclopamine, there is a possibility that cyclopamine can bind to both domains simultaneously. However, the effect on the activity of hSMO when both sites are occupied remains unclear. Based on these findings, we hypothesize that, cyclopamine can act as an agonist by binding to CRD of WT hSMO (Fig. 1a) and an antagonist by binding to TMD of WT hSMO (Fig. 1b). Additionally, we aim to discover the behavior of WT hSMO when both sites are occupied simultaneously (Fig. 1c).

**Figure 1:**
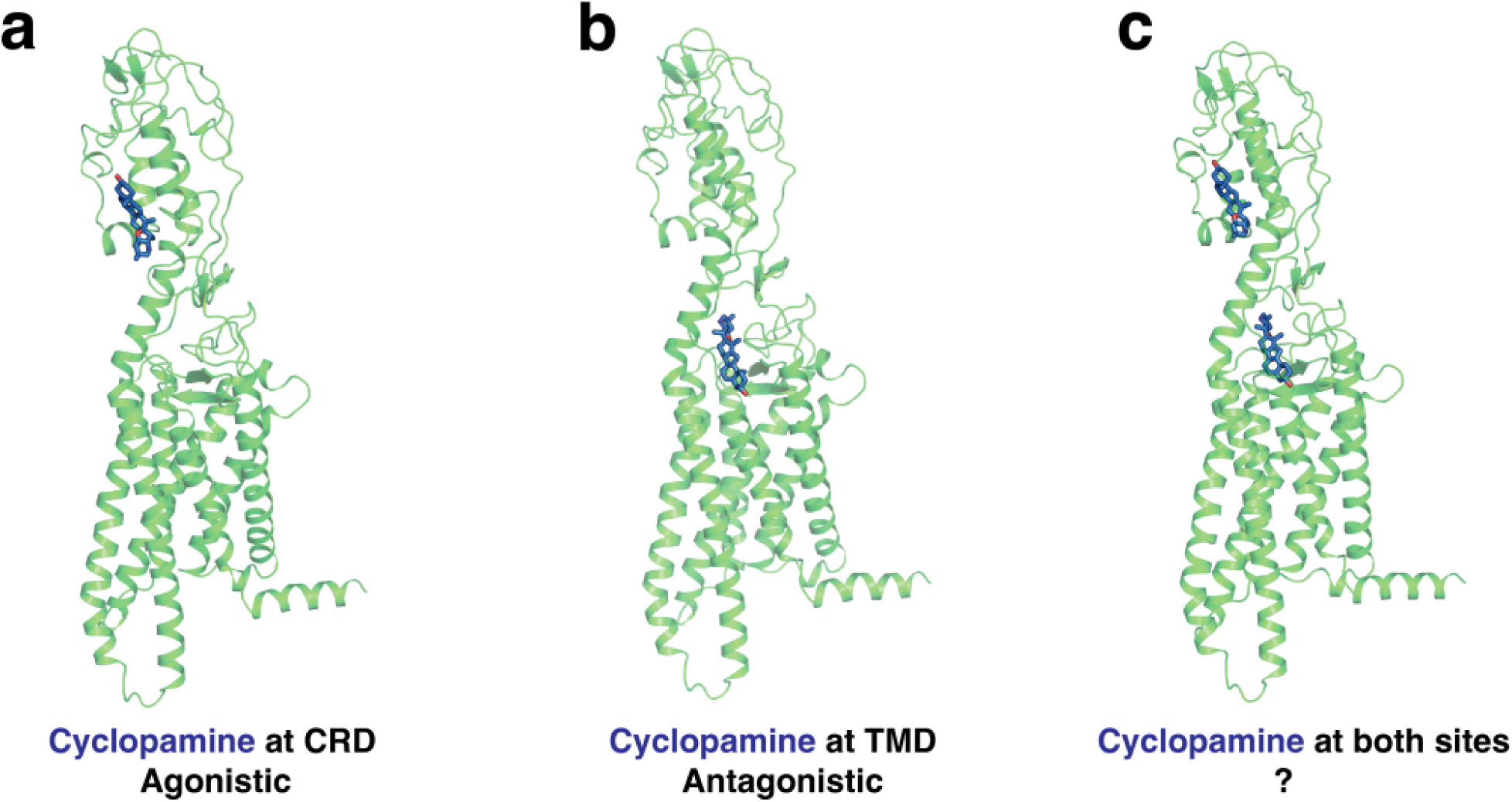
(a) Cyclopamine can bind to CRD of WT hSMO and show agonistic behavior. (b) Cyclopamine can bind to TMD of WT hSMO and show antagonistic behavior. (c) Cyclopamine can bind to both sites (CRD and TMD) simultaneously, and the net effect remains unknown.

Molecular Dynamics simulations is a powerful tool for understanding the structure-function relationship for macromolecules in atomistic detail. In a recent molecular dynamics study, the activation mechanism of human Smoothened in its ligand-free form (Apo-SMO) was explored, and the effects of the agonist SAG1 and the antagonist SANT1 on hSMO were investigated.^23^ Specifically, they analyzed the free energy barrier for Apo-SMO to undergo transition from inactive to active state. Additionally, simulations revealed that SAG1 induces an expansion of the hydrophobic tunnel inside hSMO, consistent with its cholesterol transport-like activity. Conversely, SANT1 was found to occlude the hydrophobic tunnel, thereby inhibiting hSMO activity. This data could serve as a basis for comparing and classifying cyclopamine behavior when bound to different domains of SMO. Similarly, there is another recent MD simulation study that has characterized the role of cholesterol when it binds to TMD and/or CRD of SMO and analyzed the activity of SMO for each cases.^24^

In this study, to investigate the effect of cyclopamine binding on WT hSMO, we performed MD simulations for WT hSMO bound with cyclopamine at different binding sites: at CRD, TMD, and both sites. The following names are used to refer to the simulated systems-CRD-CYC (cyclopamine bound to SMO’s CRD), TMD-CYC (cyclopamine bound to SMO’s TMD), and Dual-CYC (cyclopamine bound to SMO’s both sites). To explore the dynamics of the system, we constructed inactive and active states for each bound system as starting points for the simulations. However, long timescale associated with the activation of Smoothened precludes the use of traditional single long time-scaled MD trajectories. To overcome this limitation, we employed adaptive sampling,^25,26^ an accelerated sampling method that uses least populated frames from clusters as new starting points for further simulations. This approach facilitates efficient sampling of transitions between inactive and active states and observe the complete activation process. We have performed *∼*3 milliseconds of aggregate simulations using Markov State Model (MSMs) based adaptive sampling. However, adaptive sampling leads to statistical bias, since the methodology samples from the states to maximize the exploration of the conformational free energy landscape. One way to overcome this bias is by constructing Markov State Models,^27,28^ which divide the conformational ensemble into microstates, and estimate the rates of transitions between the states, effectively reweighing the entire ensemble. MSMs can offer precise insights into both kinetic and thermodynamic properties related to the protein dynamics. MSMs have been used extensively to characterize the conformational dynamics of GPCRs^23,29–33^ for understanding their activation mechanisms and modulation of their activity by ligands and ions.

Our results show that when cyclopamine binds to CRD of SMO, it can act as an agonist, as it showed higher active population compared to inactive at equilibrium (8% inactive, 80% active). Among all four cases (CRD-CYC, Dual-CYC, TMD-CYC, Apo-SMO), CRD-CYC showed the lowest activation barrier (2 *±* 0.2 kcal/mol), which can facilitate SMO activation. In addition, we found the tunnel to expand in the upper leaflet, to facilitate cholesterol transport and activate SMO. On the other hand, when cyclopamine binds to TMD, we show its role as an antagonist, as it showed higher inactive population compared to active at equilibrium (54% inactive, 31% active). TMD-CYC showed the highest activation barrier (4 *±* 0.2 kcal/mol), hindering the activation process. The tunnel remained blocked, which also demonstrates antagonistic behavior. In the case where cyclopamine binds to both sites, there was a slight imbalance with a higher inactive population at equilibrium (52% inactive, 41% active), suggesting weak antagonism. The activation barrier was relatively high, likely hindering the activation process (3.5 *±* 0.3 kcal/mol). Additionally, we demonstrate that cyclopamine bound at TMD has a larger effect in shrinking the tunnel, but the size of the tunnel was slightly larger than when it solely binds to TMD, which may leave a room for cholesterol transport in rare cases. Detailed residue movements upon activation showed differences in the agonistic (CRD-CYC) and antagonistic (TMD-CYC) effect of cyclopamine on SMO.

## Results and Discussion

### Binding position dependent modulation of the SMO activation process by Cyclopamine

Analyzing the equilibrium populations for the TMD-CYC, Dual-CYC, and CRD-CYC systems can give insights into the binding position dependent agonistic, antagonistic or partial agonistic behavior exhibited by cyclopamine. We employed simulation data from Apo-SMO^23^ as a reference point to discern the functional behavior of each system. To obtain equilibrium active and inactive state populations for CRD-CYC, Dual-CYC, TMD-CYC, and Apo-SMO, 57 inter-residue distances used for adaptive sampling^25^ were used as input metrics for the VampNet.^34^ VampNet uses an autoencoder architecture to perform dimensionality reduction from a set of input features. A macrostate model containing six metastable states was built using VampNets. The output of the VampNet gives the probability of a simulation frame belonging to each of the macrostates (see Methods section for details). Figure 2 shows equilibrium populations of CRD-CYC, Apo-SMO, Dual-CYC, and TMD-CYC in both the inactive and active states. The inactive to active population ratios reveal distinctive characteristics for each system: CRD-CYC (8% inactive, 80% active), Apo-SMO (49 % inactive, 44% active), Dual-CYC (52% inactive, 41% active), and TMD-CYC (54% inactive, 31% active). We find that binding of cyclopamine to SMO shifts equilibrium population in a binding position dependent manner. As shown in Figure 2, CRD-CYC has a very low inactive population as compared to active population. This suggests CRD-CYC’s agonistic behavior, as the active state is favored at equilibrium. In contrast, higher ratio of inactive to active population suggests antagonistic behavior for TMD-CYC, as it favors the inactive state. Similarly, higher ratio of inactive to active population for Dual-CYC suggests antagonism. However, its active population at equilibrium is slightly higher compared to TMD-CYC. This suggests weak antagonism of Dual-CYC.

**Figure 2:**
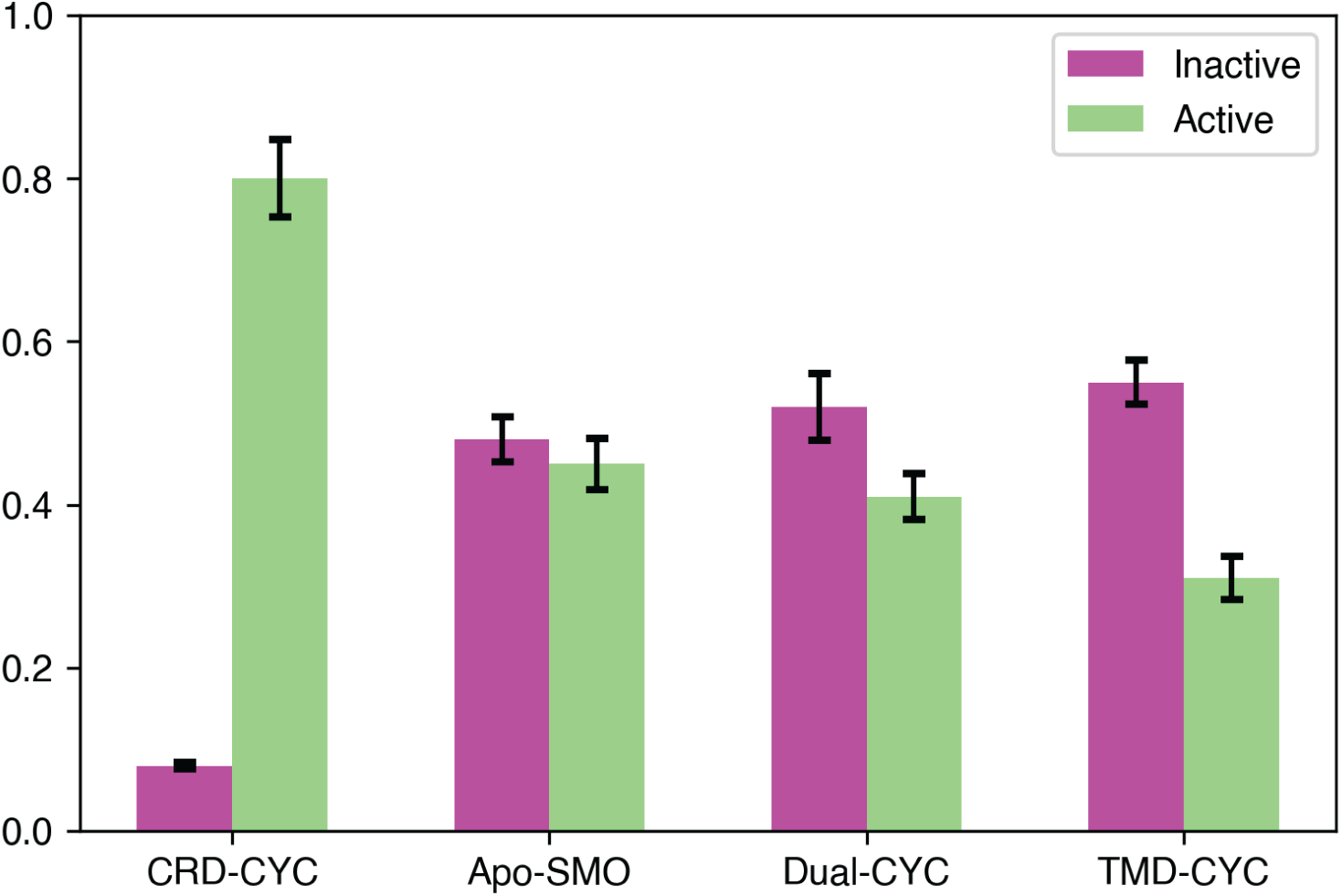
Equilibrium populations of CRD-CYC, Apo-SMO, Dual-CYC, and TMD-CYC in the inactive and active state. Inactive to active population ratio for CRD-CYC is 0.08:0.8, Apo-SMO is 0.49:0.44, Dual-CYC is 0.52:0.41, and TMD-CYC is 0.54:0.31. Error bars were computed by running the model 20 times with 80% of the data.

Breakage of E/DRY motif in Class A GPCRs, which is associated with the activation of Class A GPCRs,^1,29,35–37^ is analogous to the conserved molecular switch (W-G-M motif) in Class F receptor activation.^23^ Specifically, the outward translation of W339^3.50f^ and M449^6.30f^ and the inward translation of G422^5.65f^ have been posited to play an integral role in Class F receptor activation to accommodate *G_i_* at the intracellular end of SMO (to denote the Class F GPCR TM residues, we used modified Ballesteros-Weinstein numbering system^38^). The outward movement of M449^6.30f^ serves as a reliable indicator for the outward movement TM6,^39,40^ while W339^3.50f^ and G422^5.65f^ represent TM3 and TM5 rearrangements. To analyze the TM3-5-6 rearrangement and the overall free energy barrier associated with this rearrangement, we constructed free energy landscapes projected onto W339^3.50f^ – G422^5.65f^ (TM3-TM5 distance) and W339^3.50f^ – M449^6.30f^ (TM3-TM6 distance) for all three cases: CRD-CYC (Fig. 3a, b, Fig. S1a, Fig. S2a, b), Dual-CYC (Fig. 3c, d, Fig. S1b, Fig. S2c, d), and TMD-CYC (Fig. 3e, f, Fig. S1c, Fig. S2e, f).

**Figure 3:**
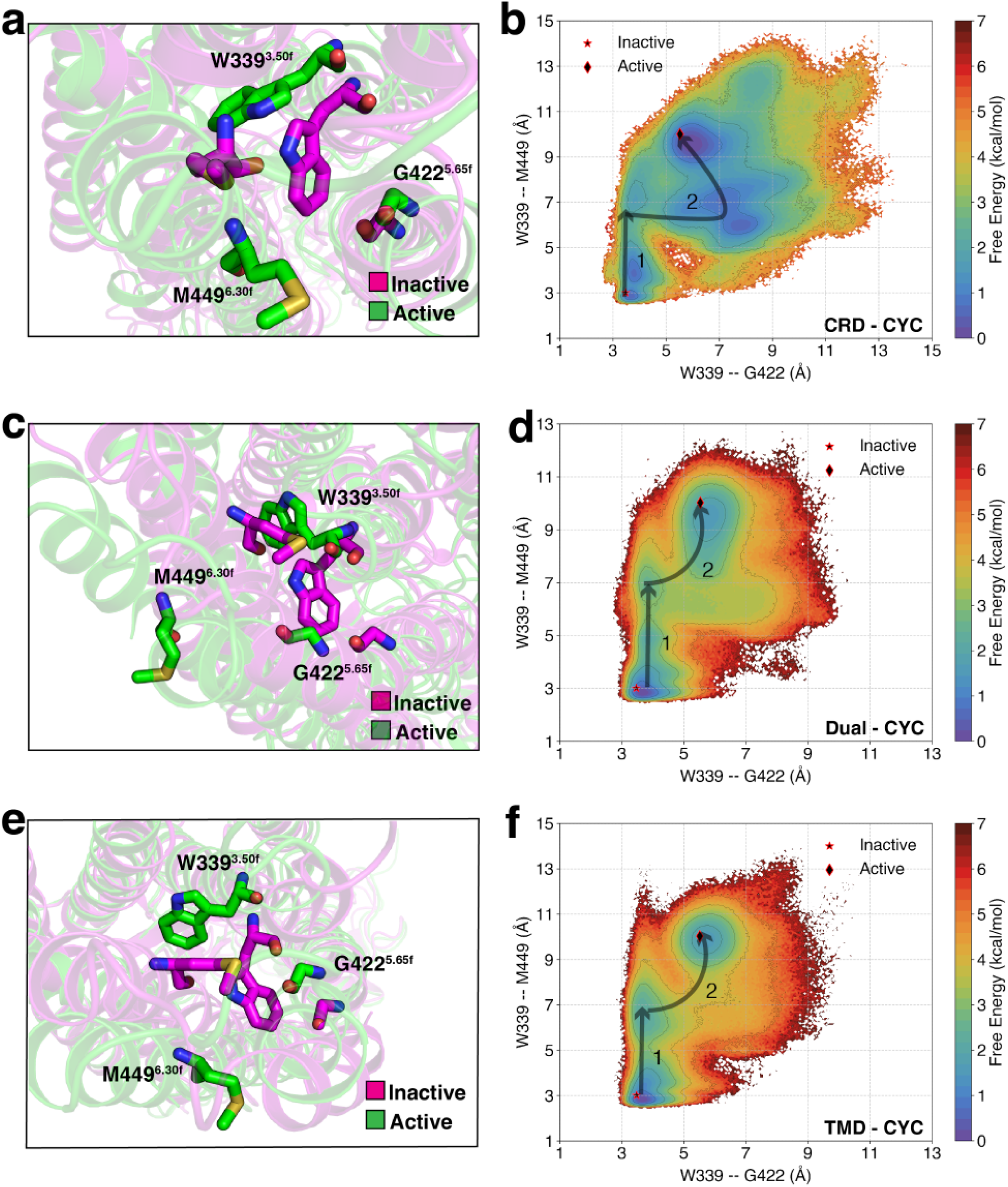
The WGM motif undergoes a rearrangement triggered by cyclopamine binding at different sites of SMO. MSM-weighted free energy landscapes projected onto W339^3.50f^ – G422^5.65f^ (TM3-TM5 distance) and W339^3.50f^ – M449^6.30f^ (TM3-TM6 distance) for (a, b) CRD-CYC, (c, d) Dual-CYC, and (e, f) TMD-CYC.

In all cases, TM6 first undergoes an outward shift of *∼* 4 Å (State 1 in Fig. 3b, d, f). After this movement, in CRD-CYC, TM3 moves outward by *∼* 4 Å, followed by a subtle TM5 rearrangement (State 2 in Fig. 3b). In contrast, TMD-CYC and Dual-CYC showed a slightly different behavior. In both cases, after the initial outward shift of TM6, TM3 experiences an outward shift of *∼* 2 Å along with a coordinated *∼* 4 Å outward shift of TM6 (State 2 in Fig. 3d, f). We find from the free energy landscapes that the overall free energy barrier for CRD-CYC is 2 *±* 0.2 kcal/mol, for Dual-CYC is 3.5 *±* 0.3 kcal/mol, and for TMD-CYC is 4 *±* 0.2 kcal/mol. The overall free energy barrier for this rearrangement in Apo-SMO showed 2.5 *±* 0.3 kcal/mol.^23^

The overall free energy barrier of CRD-CYC is lower than that of Apo-SMO. Thus, when cyclopamine binds to the CRD site, it facilitates receptor activation by reducing the activation barrier. On the other hand, the overall free energy barrier of TMD-CYC is higher than that of Apo-SMO. This shows that when cyclopamine binds to TMD site, it hinders receptor activation by increasing the activation barrier. The free energy barrier of Dual-CYC falls between CRD-CYC and TMD-CYC, but is closer to that of TMD-CYC (*∼* 0.5 kcal/mol difference). Moreover, the movement of TM3, TM5, and TM6 in Dual-CYC was similar to the movement in TMD-CYC (Fig. 3d, f). Therefore, in Dual-CYC, TMD-bound cyclopamine has a more dominant effect in the rearrangement of WGM residues and contributes to high activation barrier, to hinder the activation process.

### D-R-E network breakage facilitates smoothened activation

SAG1.5, an agonist that is known to bind to SMO’s TMD, has been observed to exert its agonistic effects through the D-R-E network, which involves residues D473^6.54f^, R400^5.43f^, and E518^7.38f^ located at the extracellular end of TMD. This network experiences disruption upon SAG1.5 binding.^41^ To examine the influence of cyclopamine on the D-R-E network during activation, especially when it binds to different domains of Smoothened (SMO), we constructed free energy landscapes projected onto R400^5.43f^ – E518^7.38f^ and W339^3.50f^ – M449^6.30f^ for CRD-CYC (Fig. 4a, b, Fig. S3a), Dual-CYC (Fig. 4c, d, Fig. S3b), and TMD-CYC (Fig. 4e, f, Fig. S3c). In this representation, R-E distance indicates forming/breaking of the salt bridge between the basic residue arginine (R) and the acidic residue glutamic acid (E). Furthermore, we focused on W339^3.50f^ – M449^6.30f^ as it reflects the intracellular movement between TM3-TM6, the key indicator of the activation process of SMO (Fig. S2b, d, f). Hence, constructing the free energy landscapes can help us analyze how the D-R-E network undergoes changes in response to the activation of SMO.

**Figure 4:**
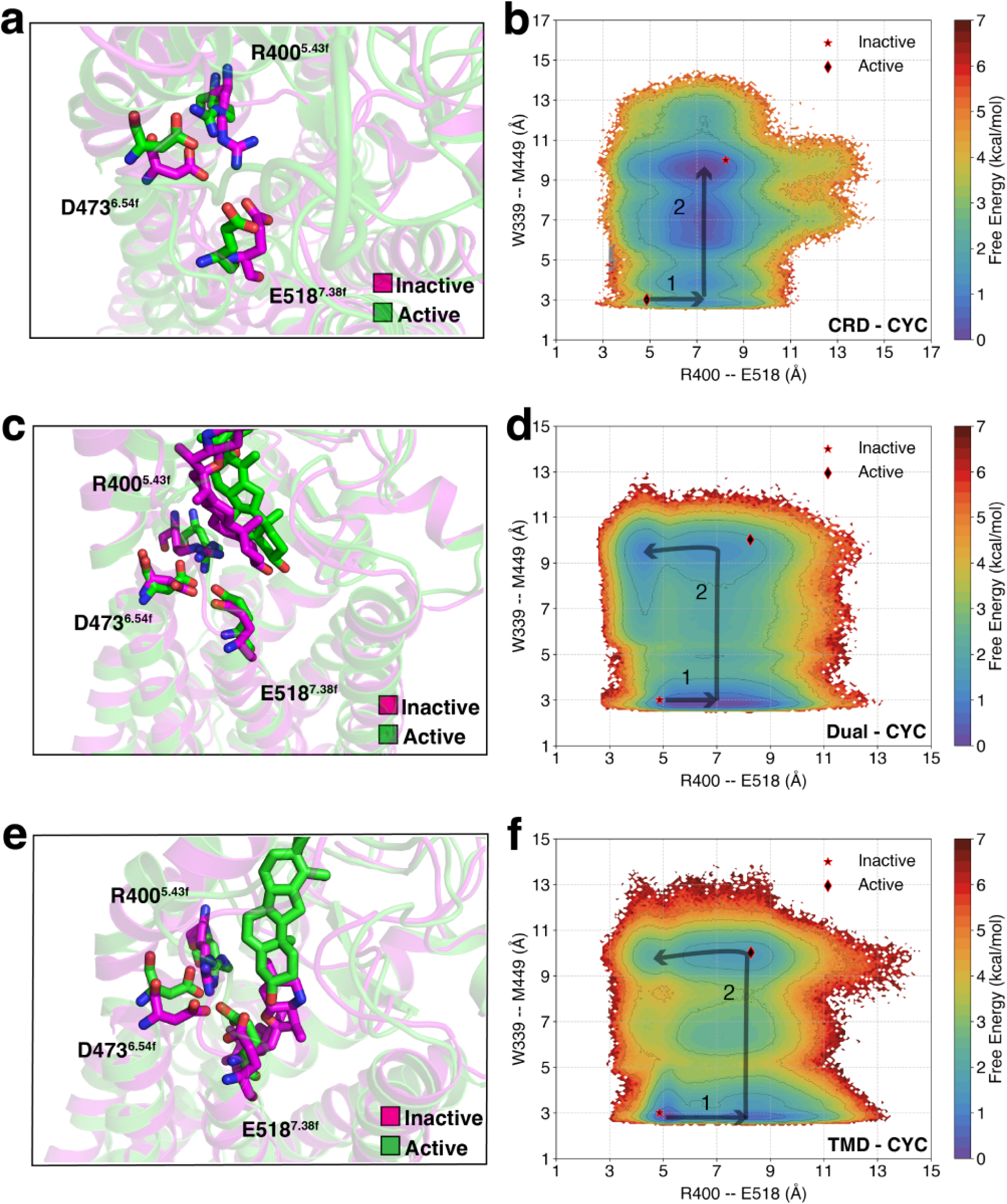
D-R-E network undergoes a rearrangement triggered by cyclopamine binding at different sites of SMO. MSM-weighted free energy landscapes projected onto R400^5.43f^ – E518^7.38f^ and W339^3.50f^ – M449^6.30f^ (TM3-TM6 distance) for (a, b) CRD-CYC, (c, d) Dual-CYC, and (e, f) TMD-CYC.

In CRD-CYC, a clear D-R-E network breakage is observed upon SMO activation. Following the breakage of the salt bridge between R400^5.43f^ and E518^7.38f^ from *∼* 5 Å to *∼* 7 Å (State 1 in Fig. 4b), an increase in TM3-TM6 distance is observed (State 2 in Fig. 4b). The Dual-CYC and TMD-CYC systems also follow the initial perturbation, where the R400^5.43f^ – E518^7.38f^ distance extends from *∼* 5 Å to *∼* 7 Å (State 1 in 4d, f). Upon the increase of TM3-TM6 distance, in the active state, it could remain broken or close back to its initial state (State 2 in Fig. 4d, f). This shows that the D-R-E network breakage can facilitate SMO activation and is independent of the binding sites of cyclopamine. In addition, Dual-CYC showed similar residue movements to that of TMD-CYC. This shows that cyclopamine bound at TMD has a more dominant effect in D-R-E network movement upon activation.

### Structural dynamics and kinetic insights into cyclopamine-mediated activation

We used time-lagged independent component analysis (tICA) method to identify the linear combination of inter-residue distances that exhibit slowest decorrelation time. The first two time-lagged independent components (tICs) represent the two slowest processes observed in simulations. In our case, tIC1 correlates with the activation process from inactive state to active state for all cases. We projected the simulation data along the first two tICs for CRD-CYC (Fig. 5a), Dual-CYC (Fig. 5c), and TMD-CYC (Fig. 5e). In CRD-CYC, we identified distinct minima corresponding to inactive and active states. However, we observed a spread of density along tIC2 (Fig. 5a), which could be attributed to SMO’s CRD being decoupled from TMD upon CYC binding (Fig. S5a, b). This further explains the CRD’s role in SMO activation, as previous studies have shown that SMO’s CRD suppresses its basal activity. ^42^ Hence, when the decoupling happens in CRD-CYC, the suppressive effect of the CRD on SMO is relieved and the activation barrier is reduced.

**Figure 5:**
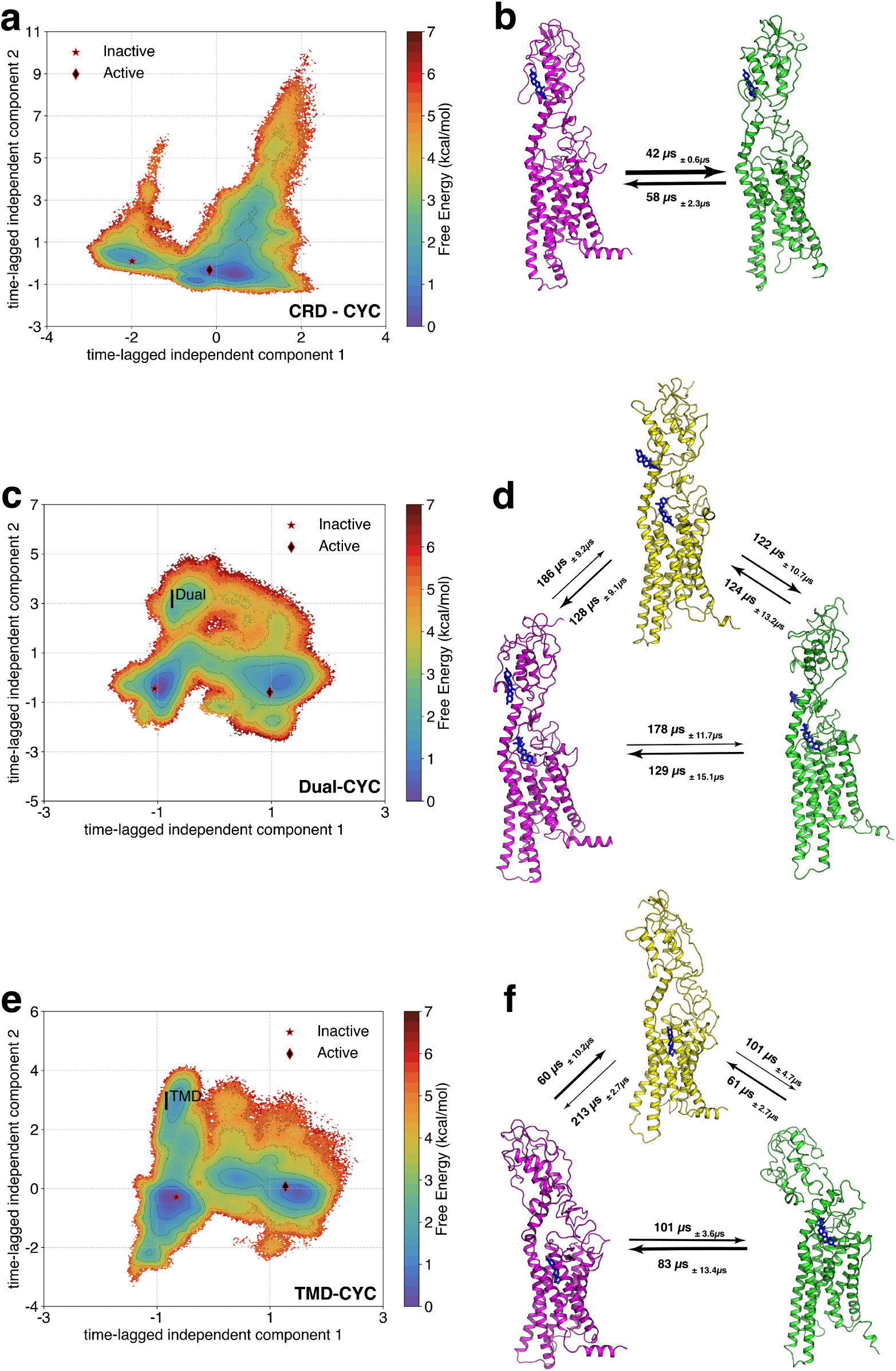
tIC plots and the corresponding MFPT analysis for (a,b) CRD-CYC, (c,d) Dual-CYC, and (e,f) TMD-CYC. Inactive (magenta) and active (green) states were identified in all cases. Intermediate states (yellow) were identified in Dual-CYC (I^Dual^) and TMD-CYC (I^TMD^).

On the other hand, in Dual-CYC and TMD-CYC, we identified intermediate states (I^Dual^ and I^TMD^) along with inactive and active states from each minima (Fig. 5c, e). To ascertain the specificity of these intermediate states within their respective systems, we projected simulation data onto the tICA space of the other two systems (Fig. S6). When Dual-CYC data was projected onto the tIC space of TMD-CYC, I^TMD^ remained evident (Fig. S6f). This indicates that I^TMD^ is not exclusive and is present in both Dual-CYC and TMD-CYC. Conversely, when TMD-CYC data was projected onto the tIC space of Dual-CYC, I^Dual^ was not observed (Fig. S6d). This highlights the distinctiveness of I^Dual^. To quantify the uniqueness of I^Dual^, we computed the Kullback-Liebler Divergence (KL Divergence) for I^Dual^ with the active and inactive states as a reference (explained in Methods). Conformational changes occur in ECL1 and TM1 during the transition from the inactive state to I^Dual^ (Fig. S5c), as these regions showed the highest K-L divergence. In addition, conformational changes occur in CRD during the transition from I^Dual^ to the active state (Fig. S5d). The conformational change in cyclopamine-bound CRD effectively explains Dual-CYC’s transition to the active state, as the suppressive effect of CRD on SMO may partially be relieved due to cyclopamine binding to CRD. ^42^ In TMD-CYC, we found conformational changes occurring in the loops surrounding G80^CRD^ during both the transition from the inactive state to I^TMD^ (Fig. S5e) and the transition from I^TMD^ to the active state (Fig. S5f). Overall, the conformational change in TMD-CYC is most restricted, indicating its preference to remain in inactive conformation.

The Mean First Passage Time (MFPT) analysis is an important tool for assessing the time required for a system to transition between different states within the MSM framework.^43^ We applied Transition Path Theory^44,45^ to our constructed MSM, to compute the transition fluxes between these states and establish timescales associated with the activation for CRD-CYC (Fig. 5b), Dual-CYC (Fig. 5d), and TMD-CYC (Fig. 5f). The observed differences in transition times between the inactive and active states in CRD-CYC, Dual-CYC, and TMD-CYC are related to their respective kinetic properties.

In CRD-CYC, the transition from the inactive to active state occurs approximately 1.4 times faster than the reverse process (Fig. 5b). Additionally, the pathway from the inactive state to the active state doesn’t show any additional metastable states. In contrast, Dual-CYC and TMD-CYC show additional intermediate states, which lead to a higher kinetic barrier. This increases the timescales to achieve activation in these cases, as the timescales depend on the total flux between these states. The overall transition timescales are *<*100 *µ*s for CRD-CYC, compared to *>*100 *µ*s for TMD-CYC and Dual-CYC. MFPT analysis in CRD-CYC suggests a favorable kinetic pathway for activation, likely attributed to a lower activation energy barrier and more efficient conformational changes leading to the active state. On the other hand, transition flux of Dual-CYC and TMD-CYC indicates a kinetic preference towards the inactive state and imply a more complex or energetically demanding process for activation, involving additional steps for structural rearrangements that require a longer time for activation.

### Cyclopamine at CRD expands SMO’s hydrophobic tunnel

A unique feature of SMO is the presence of an internal tunnel, which plays an important role in facilitating the transfer of cholesterol from the cell membrane to the binding site in CRD.^39,40,42,46,47^ Composed of hydrophobic residues, the tunnel starts from W339^3.50f^, extends across approximately seven transmembrane helical turns, and terminates at the D-R-E network (D473^6.54f^, R400^5.43f^, E518^7.38f^). SMO antagonists (SANT1, AntaXV, and LY2940680) are known to bind deep within this tunnel and obstruct the tunnel within SMO. On the other hand, SMO agonists (SAG) can bind outside of the tunnel and activate SMO and expands its tunnel to transport cholesterol. MD simulation of Apo-SMO, SMO bound to SANT1 (antagonist), and SMO bound to SAG (agonist) further corroborate this hypothesis, by demonstrating the clear expansion of the tunnel when SMO is bound to SAG.^23^

To investigate the effect of cyclopamine on tunnel expansion of SMO, we conducted the tunnel analyses in CRD-CYC, Dual-CYC, and TMD-CYC using HOLE program.^48^ In CRD-CYC, we observed an expansion of the tunnel between z = 0 and z = 20 Å, corresponding to the upper leaflet of the membrane (Fig. 6a, b). We computed free energy difference between the different systems to ascertain the cyclopamine’s binding position dependent effects on tunnel expansion (Fig. S8). Compared to Dual-CYC and TMD-CYC, CRD-CYC clearly showed the expansion in the upper leaflet of the membrane (Fig. S8a, b). This agrees with the expansion of tunnel for SAG bound SMO at the upper leaflet of the membrane.^23^ This suggests that the presence of cyclopamine bound at CRD induces a relative enlargement of the tunnel. The exact location of the tunnel in the upper leaflet opening corresponded to the region between TM5 and TM6 (Fig. S7a, b). This agrees with the a recent study that observed hydrophobic tunnel opening of active SMO at TM5 and TM6.^46^

**Figure 6:**
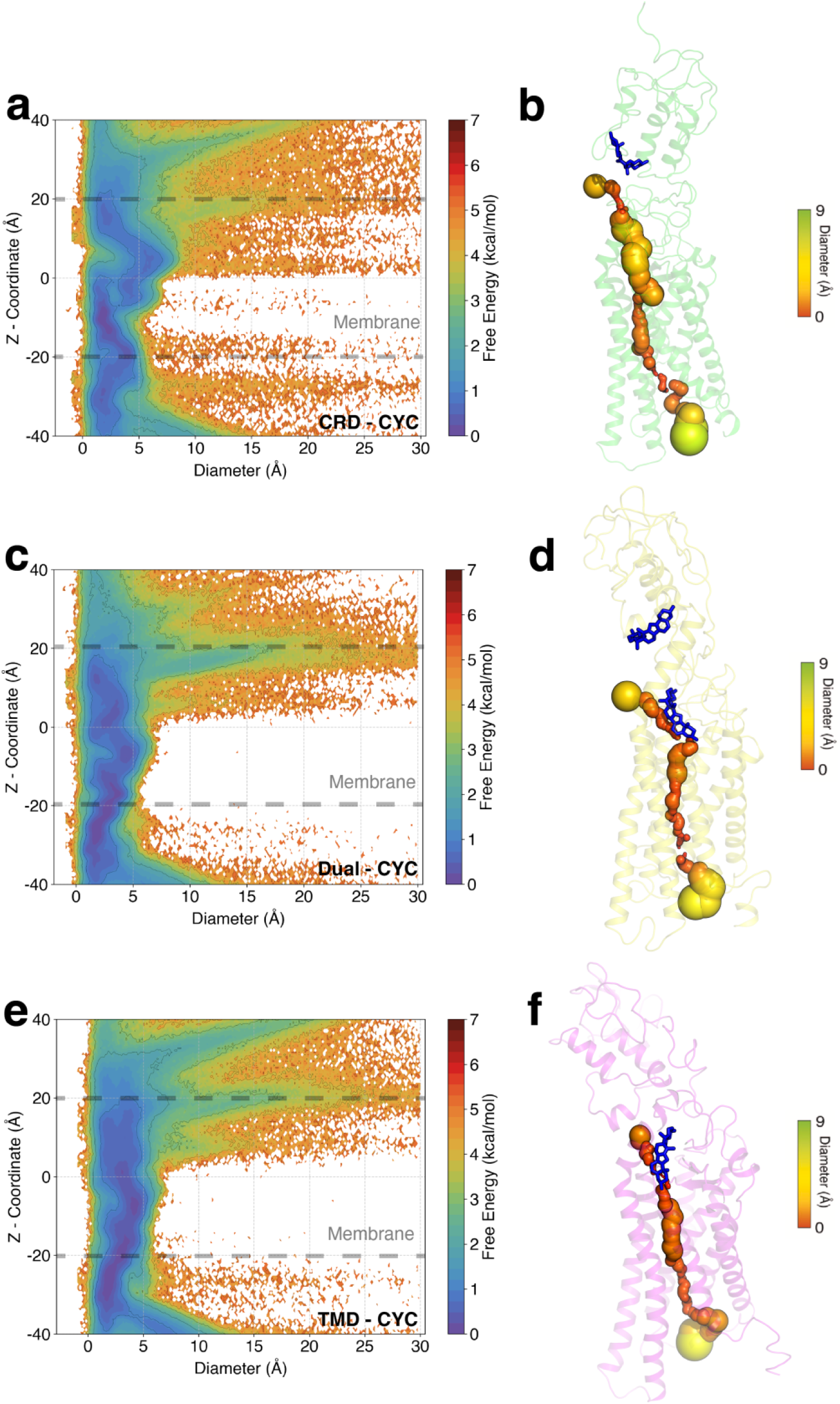
Tunnel diameter plots for the SMO. (a) Free energy plot of the tunnel diameter along the z-coordinate for CRD-CYC, (c) for Dual-CYC, and (e) TMD-CYC. (b), (d), (f)-representative SMO with internal tunnels. CRD-CYC shows a clear expansion of the tunnel in the upper leaflet compared to TMD-CYC and Dual-CYC.

In TMD-CYC, the tunnel remained obstructed (Fig. 6e, f). This indicates that cyclopamine binds within the core of the SMO tunnel, effectively impeding the transport of cholesterol and blocking SMO activity. In Dual-CYC, the tunnel is also majorly blocked (Fig. 6c, d) but it is relatively larger as compared to TMD-CYC (Fig. S8c), though this enlargement is not as significant as what was observed in CRD-CYC (Fig. S8a).

### Identification of key residues that balance agonistic and antagonistic behavior during SMO activation

To delineate specific residue movements for CRD-CYC, Dual-CYC, and TMD-CYC during activation, a multi-class Random Forest Classifier was used. The goal of the classifier was to identify features that distinguish the three ensembles. The input to the classifier consisted of the 57 distances used for adaptive sampling,^25^ 494 *ψ* backbone dihedrals to characterize backbone movements, and 234 *χ*_2_ dihedrals to characterize sidechain movements. 5-fold cross-validation and hyperparameter optimization was applied to the model to identify the top 20 differentiating features (explained in Methods). After identifying the unique residue movements in CRD-CYC and TMD-CYC, we further classified Dual-CYC’s behavior based on these distinct patterns. Our findings unveiled that Dual-CYC exhibited a heterogeneous behavior in terms of residue movements, which lies between the agonistic and antagonistic behavior of CRD-CYC and TMD-CYC.

In CRD-CYC, the *χ*_2_ dihedral angle at Y472^6.53f^, which is a conserved residue across all Class F GPCRs, remained stable (Fig. 7a, b). In contrast, we observed a 180° rotation of this angle in TMD-CYC (Fig. 7e, f). In Dual-CYC, we also observed a 180° rotation of this angle, showing antagonistic behavior (Fig. 7c, d). This may be attributed to cyclopamine bound at TMD exerting a dominant effect on rotating this angle movement. In CRD-CYC, we also observed a 40 ° partial rotation of *ψ* dihedral angle between F526^7.46f^ and G527^7.46f^ (Fig. S9a), which are partially conserved across Class F GPCRs. In contrast, this angle remained stable in TMD-CYC (Fig. S9c). In Dual-CYC, this angle also remained stable (Fig. S9b). This shows that cyclopamine bound at TMD exerts a dominant effect in restricting this angle movement. These distinctive features characterize antagonism within Dual-CYC.

**Figure 7:**
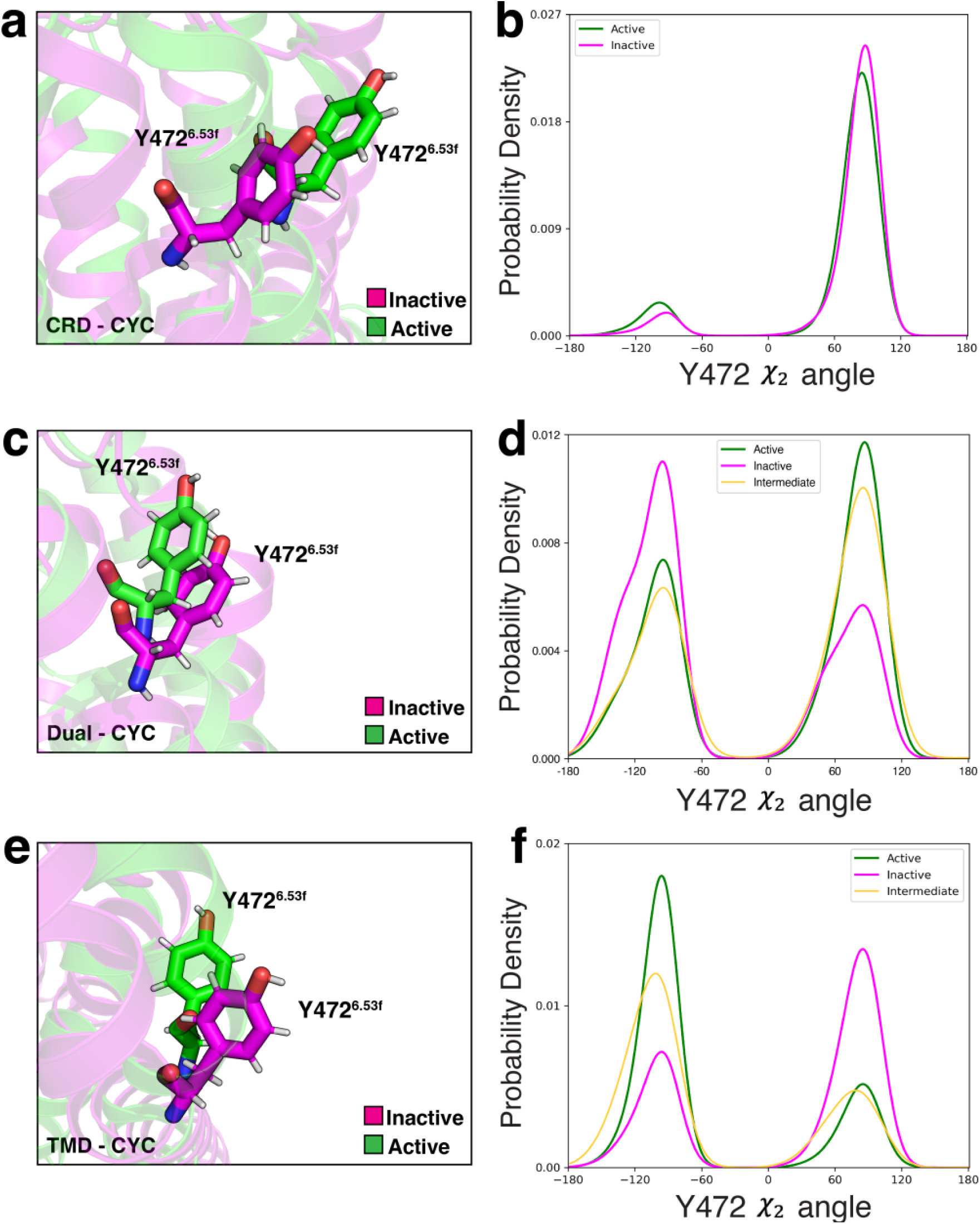
Snapshots and probability density plots of *χ*_2_ dihedral angle rotation at Y472^6.53f^ in (a,b) CRD-CYC, (c,d) Dual-CYC, (e,f) TMD-CYC.

Agonistic attributes were also identified in Dual-CYC. We identified 180° rotation in the *ψ* dihedral angle between T90^CRD^ and L91^CRD^ in CRD-CYC (Fig. S9d), contrasting with its steady state in TMD-CYC (Fig. S9f). In Dual-CYC, we observed 180° rotation of this angle (Fig. S9e), and this may be attributed to cyclopamine bound at CRD exerting a dominant effect on this angle rotation. The *ψ* dihedral angle between the conserved residue C213^CRD^ and G214^CRD^ remained stable in CRD-CYC (Fig. S9g), while a 220° rotation was observed in TMD-CYC (Fig. S9i). In Dual-CYC, we observed this angle to also remain stable (Fig. S9h). This may be attributed to cyclopamine bound at CRD exerting a dominant effect on locking this angle movement. Lastly, we identified 180° rotation in the *χ*_2_ dihedral angle at the conserved residue W535^7.55f^ in CRD-CYC (Fig. S9j), leading to the breakage of the *π*-cation interaction between W535^7.55f^ and R451^6.32f^. In contrast, this angle remained static in TMD-CYC (Fig. S9l). In Dual-CYC, this angle was rotated by 180° leading to the breakage of the *π*-cation interaction (Fig. S9k). These observations collectively suggest that Dual-CYC exhibits a combination of both agonistic and antagonistic features in terms of residue movements.

## Conclusions

Our study provides a detailed understanding of hSMO activation when bound to cyclopamine at distinct domains-CRD, TMD, and both domains. The simulation data from Apo-SMO was employed as a reference point,^23^ to discern the relative behavior for each case. Equilibrium population analysis showed that CRD-CYC favors the active state (8% inactive, 80% active), suggesting agonistic behavior. In contrast, TMD-CYC favors the inactive state (54% inactive, 31% active), indicating antagonistic behavior. In the case of Dual-CYC, there was a relative balance between the inactive and active populations at equilibrium (52% inactive, 41% active). The slightly higher inactive population compared to active population shows weak antagonism.

We found the breakage of the D-R-E network facilitates hSMO activation upon cyclopamine binding for all three cases. In addition, we demonstrated that all three cases can undergo activation through the rearrangement of an intracellular structural motif known as the W-G-M motif, a conserved feature in Class F GPCRs. Among all four cases (CRD-CYC, Apo-SMO, Dual-CYC, and TMD-CYC), CRD-CYC showed the lowest overall free energy barrier (activation barrier) for this rearrangement, with 2 *±* 0.2 kcal/mol, which can facilitate SMO activation. In contrast, TMD-CYC showed the highest overall free energy barrier, with 4 *±* 0.2 kcal/mol, hindering the activation process. Dual-CYC also showed high overall free energy barrier, with 3.5 *±* 0.3 kcal/mol, hindering the activation process. Along with the analysis of activation energy barrier, we showed that the transition pathway theory analysis demonstrates the kinetically favorable pathway in CRD-CYC, likely attributed to low activation energy barrier. In contrast, Dual-CYC and TMD-CYC indicated its kinetic preference towards the inactive state, likely attributed to high activation energy barrier. The tunneling analysis also provides clues to analyze the SMO activity for the three cases. CRD-CYC shows agonistic character, as we observe the expansion of the hydrophobic tunnel in CRD-CYC in the upper leaflet to facilitate the cholesterol transport, which can lead to the activation of SMO. In contrast, in TMD-CYC, the tunnel remained obstructed, impeding the cholesterol transport, inhibiting SMO. In Dual-CYC, the tunnel was slightly larger than TMD-CYC but smaller than CRD-CYC. This could be due to cyclopamine bound at CRD inducing a small enlargement on the tunnel to transport cholesterol on rare cases.

In CRD-CYC, we found the major conformation changes occurring in CRD during SMO activation, which may be attributed to cyclopamine relieving the suppressive effect of CRD on SMO.^42^ On the other hand, TMD-CYC shows the most restricted conformational changes upon activation, occurring primarily in G80^CRD^. In Dual-CYC, we find conformational changes occurring in TM1 and ECL1 during transition from inactive to I^Daul^, and in CRD during the transition from I^Daul^ to active state. This suggests that cyclopamine bound at CRD can play a role and affect the latter transition in Dual-CYC. Remarkably, a more detailed examination of residue movements in Dual-CYC showed a balance between agonistic and antagonistic behaviors of CRD-CYC and TMD-CYC.

Throughout these analyses, we collectively demonstrate CRD-CYC’s agonistic behavior, TMD-CYC’s antagonistic behavior, and Dual-CYC’s weak antagonistic behavior. Our study provides crucial insights into the dynamics of hSMO activation with cyclopamine binding to distinct domains. In particular, the identification of such unique residue movements within Dual-CYC can open up new possibilities for drug development. These findings will hold potential implications for targeted therapeutic interventions in disorders associated with Hh signaling pathways.

## Methods

### Molecular dynamics simulations

#### Simulation setup

To construct inactive CRD-CYC, we aligned xSMO (Xenopus Smoothened) bound with cyclopamine at both sites (PDB ID: 6D32^47^) and inactive hSMO (PDB ID: 5L7D^42^), and then removed xSMO, cyclopamine bound at TMD, and stabilizing antibodies. To construct active CRD-CYC, we aligned xSMO (Xenopus Smoothened) bound with cyclopamine at both sites (PDB ID: 6D32^47^) and active hSMO (PDB ID: 6XBL^40^), and then removed xSMO, cyclopamine bound at TMD, and stabilizing antibodies. To construct inactive TMD-CYC, we aligned cyclopamine bound hSMO lacking CRD (hSMOΔCRD) (PDB ID: 409R ^49^) and inactive hSMO (PDB ID: 5L7D^42^), and then removed hSMOΔCRD and stabilizing antibodies. To construct active TMD-CYC, we aligned cyclopamine bound hSMO lacking CRD (hSMOΔCRD) (PDB ID: 409R^49^) and active hSMO (PDB ID: 6XBL^40^), and then removed hSMOΔCRD and stabilizing antibodies. To construct inactive Dual-CYC, we aligned inactive TMD-CYC and active CRD-CYC, and then removed active hSMO structure. To construct active Dual-CYC, we aligned inactive TMD-CYC and active CRD-CYC, and then removed inactive hSMO structure. Alignment, removal of ligands, and stabilizing antibodies were performed using PYMOL.^50^ For each system, we used MODELLER^51^ to model missing residues (Table S1). In all SMO systems, E518 and H227 were protonated to match the physiological conditions.^23^ We closed the terminal residues using neutral terminal caps acetyl (ACE) for the N-terminus and methylamide (NME) for the C-terminus. The proteins were embedded in a membrane bilayer using CHARMM-GUI.^52^ ^53^ CHARMM36 force field was used to characterize the atomic interactions.^54^ ^55^ The membrane bilayer was formed using a lipid composition inspired by the lipid makeup of the cerebellum in mice brain^56^ (75% 1-Palymitoyl-2-oleoylphosphatidylcholine (POPC), 21% cholesterol, 4% sphingomyelin) (Table S2). The system was hydrated using TIP3P water^57^ and supplemented with 150mM NaCl. The total number of atoms for inactive TMD-CYC, active TMD-CYC, inactive CRD-CYC, active CRD-CYC, inactive Dual-CYC, active Dual-CYC were 106056 atoms, 104357 atoms, 105894 atoms, 104552 atoms, 105995 atoms, and 104434 atoms, with box sizes 86 *×* 86 *×* 154 Å^3^, 86 *×* 86 *×* 152 Å^3^, 86 *×* 86 *×* 154 Å^3^, 86 *×* 86 *×* 152 Å^3^, 86 *×* 86 *×* 154 Å^3^, and 86 *×* 86 *×* 152 Å^3^. The mass of non-protein hydrogens was repartitioned to 3.024 Da, to enable simulations with a longer timestep of 4 femtoseconds (fs).

#### Pre-production MD

AMBER18 was used for biomolecular simulations.^58^ ^59^ ^60^ ^61^ Pre-production MD involves multiple steps. Initially, the system was minimized for 1000 steps, using steepest descent method. The system was further minimized for 14000 steps, constraining hydrogen-containing bonds, using SHAKE algorithm.^62^ The system was then heated from 0 K to 310 K under NVT ensemble for 5 ns. The system was then equilibrated for 310 K and 1 bar for 5 ns under NPT ensemble. This was followed by equilibration for 40 ns.

#### Production MD

GPU-accelerated pmemd.cuda package from AMBER18^58^ was used for the production MD simulations. The integrator timestep used in the production MD simulations was 4 fs. Periodic boundary conditions were used. Langevin thermostat^63^ was used to maintain the temperature to mimick a constant temperature environment. The pressure of the systems was set to 1 bar and maintained using the Monte Carlo barostat. The particle mesh Ewald (PME) method^64^ was used to compute long-range electrostatic interactions. The SHAKE algorithm^62^ was used to restrain the hydrogen bonds. The cutoff for non-bonded interactions (e.g., van der Waals interactions) was set to 10 Å. Frames of the simulation were saved every 25,000 steps, giving a frame rate of 100 ps between each frame.

#### Adaptive Sampling and MSM construction

To overcome the limits of traditional MD simulation, least-count based adaptive sampling method^25,26^ was performed using pyEMMA python library.^65^ Recently, a variety of machine learning techniques have been integrated with machine learning to develop improved sampling schemes.^66–69^ However, the least count based sampling still provides the simplest framework for adaptive sampling especially for the cases where reaction coordinates for the conformational change are not available *a priroi*. The least sampled conformations generated from adaptive sampling were used as starting points for subsequent rounds of simulations. 57 pairs of distances, which is calculated from Δ residue-residue contact score (RRCS),^23,36^ were used as adaptive sampling metrics to simulate the full transitions from inactive to active state (Table S3, S4, S5, S6). Prior to MSM construction, time-lagged Independent Component Analysis (tICA) was performed to reduce the high dimensionality of data^70^ (Fig. S10, S11, S12). To identify the optimal parameters for each system, including the number of clusters and tICA components, the dimensionality of the data was reduced to five distinct tICA dimensions (3, 5, 8, 11, 14). For each configuration, the data was clustered with varying numbers of clusters using k-means clustering. VAMP2 scores were then computed for each case, and the ideal number of clusters and tICA components were determined based on the highest VAMP2 score and the convergence of the implied timescales concerning the MSM lagtime. The resulting parameters for CRD-CYC were 150 clusters and 11 tICA components (Fig. S13a, b), Dual-CYC were 400 clusters and 14 tICA components (Fig. S14a, b), and TMD-CYC were 150 clusters with 11 tICA components (Fig. S15a, b). The chosen MSM lagtime for all three systems was 30 ns. To validate the MSM, the Chapman-Kolmogorov test was performed on five macrostates using the pyEMMA python library^65^ (Fig. S16, S17, S18), and Raw counts versus MSM population were plotted for each system (Fig. S19).

### Analyzing the major structural changes of intermediate states

To analyze the major structural changes of intermediate states, closest heavy carbon atom distances for each residue pair was calculated on 50000 frames extracted from each metastable states. The distribution of distances between specific pairs of residues was then compared to the corresponding distributions in different metastable states using Kullback-Leibler (K-L) divergence analysis.^33,71^

### Differentiating between the different ensembles using multi-class Random Forest Classifier

To differentiate the residue movements in CRD-CYC, Dual-CYC, and TMD-CYC, we employed metrics that are agnostic to the choice of an expert and provide insights into the biophysical reasons governing the different behavior of each system. We employed Random-Forest classifier, which uses metrics calculated from the entire ensemble of systems, along with the labels used to differentiate between the 3 ensembles. The input metrics used were the 57 distances (Table S3), the 494 *ψ* dihedrals, and the 234 *χ*_2_ dihedrals. The output label to be predicted by the model was either 1, 2 or 3 (denoting CRD-CYC, Dual-CYC and TMD-CYC, respectively). 5-fold cross-validation was performed in addition to a GridSearch to optimize the hyperparameters for the model (max depth=20 and n estimators=100) (Fig. S20). Top 20 features that were considered important for differentiating each system were identified. The entire implementation was done using sklearn’s *sklearn.ensemble.RandomForestClassifier* module.^72^

### Identification of macrostates using VAMPnets for equilibrium population calculations

For estimating the population of the active and inactive states in each ensemble, VAMPnets,^34^ a deep learning-based method was used. VAMPnets consists of two lobes, each consisting of fully connected layers. One lobe takes the instantaneous dataset (*x_t_*) while the other uses a time-lagged dataset (*x_t_*_+_*_τ_*) to conduct a non-linear dimension reduction. The features used for MSM construction were used as input dataset. The optimization is achieved by maximizing the VAMP2-score. The lagtime (*τ*) is chosen to ensure complete de-correlation between the chosen datapoints. The lagtime chosen for VAMPnet construction was the same as the MSM lagtime, 30 ns. The output of VAMPnets is the probability of a particular frame belonging to a certain macrostate-which can then be used to assign macrostate to each frame. The probabilities were also calculated for the inactive and active starting frames, and all frames within the same macrostate as the inactive/active starting frames were considered inactive/active. The populations of the inactive/active macrostates were then calculated based on the number of frames present in that macrostate, and reweighed using the MSM probabilities. Accordingly, 6 macrostates were chosen for each ensemble. To train the VAMPnets, the PyTorch deep learning library^73^ was used.

## Trajectory analysis and visualization

Trajectory processing tasks were performed using cpptraj.^74^ Visualization and image rendering were carried out using VMD^75^ and PyMOL. MDTraj^76^ was utilized to compute distances, *ψ* and *χ*_2_ dihedrals. Matplotlib^77^ was used for plot creation. Numerical computations were aided with Numpy.^78^ Tunnel radii calculations for each z coordinate of all systems were performed using HOLE program.^48^

## Acknowledgement

Authors thank Soumajit Dutta and Austin Weigle of the Shukla Group at University of Illinois for providing valuable insights in this study. The authors thank Folding@Home distributed computing system for providing computational resource for this work. This study is supported by NIH grant R35GM142745.

## Supporting Information Available

Codes used for analysis are available at github: https://github.com/ShuklaGroup/SMO_ CYC. Trajectories and parameter files are available on Box: https://uofi.box.com/s/ 4g3xmumfmesb68y7tb0fn8wvhvycylrf.

