## Supplementary Information for "Binding Position Dependent Modulation of Smoothened Activity by Cyclopamine"

### Supplementary Information
### Binding Position Dependent Modulation of Smoothened Activity by Cyclopamine

Kihong Kim,[†] Prateek Bansal,[†] and Diwakar Shukla[*,†,‡,¶,§]

†*Department of Chemical and Biomolecular Engineering, University of Illinois at Urbana-Champaign, Urbana, IL, 61801, United States*

‡*Department of Bioengineering, University of Illinois at Urbana-Champaign, Urbana, IL, 61801, United States*

¶*Center for Biophysics and Quantitative Biology, University of Illinois at Urbana-Champaign, Urbana, IL, 61801, United States*

§*Cancer Center at Illinois, University of Illinois at Urbana-Champaign, Urbana, IL, 61801, United States*

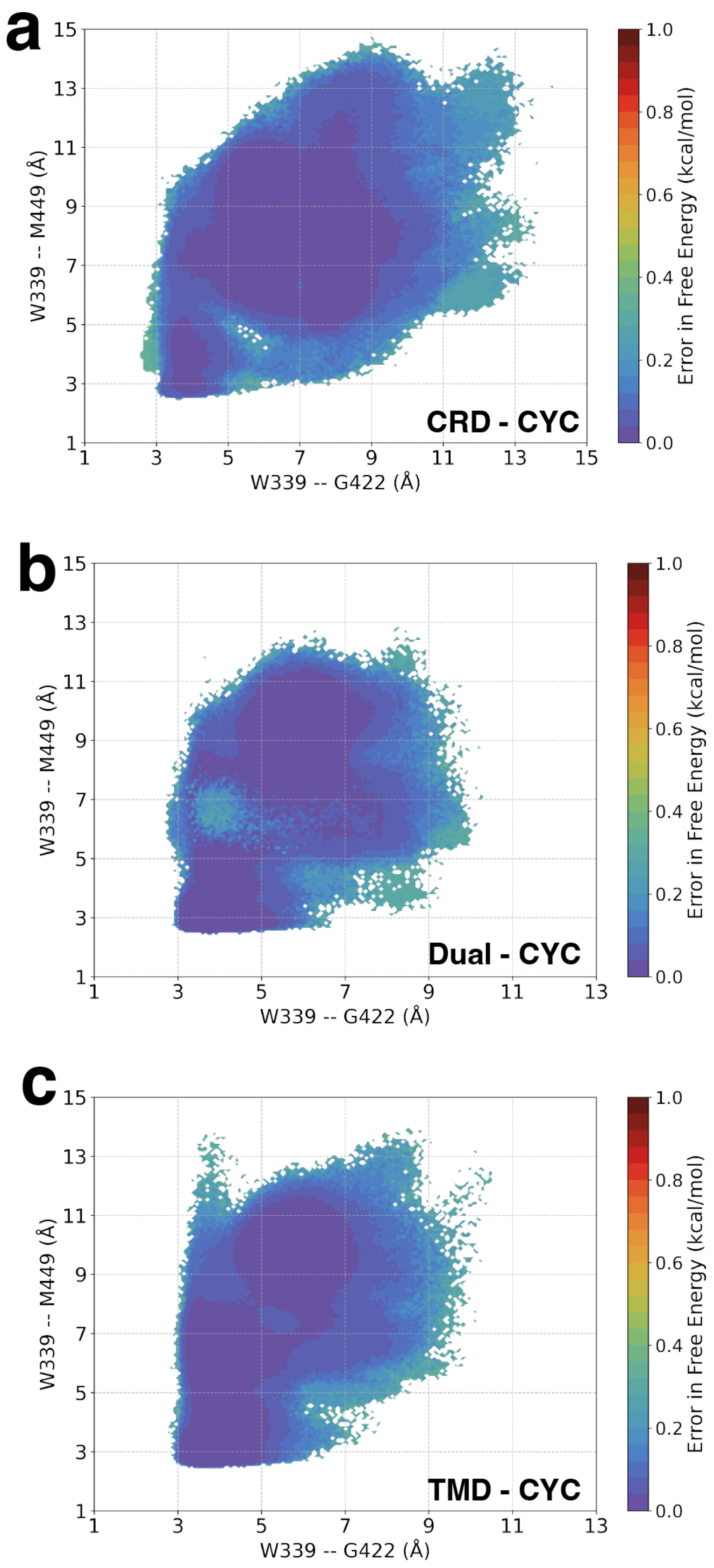


Figure S1: Errors in Figure 2 were calculated by using bootstrapping. 80% of the data was used in 200 iterations.

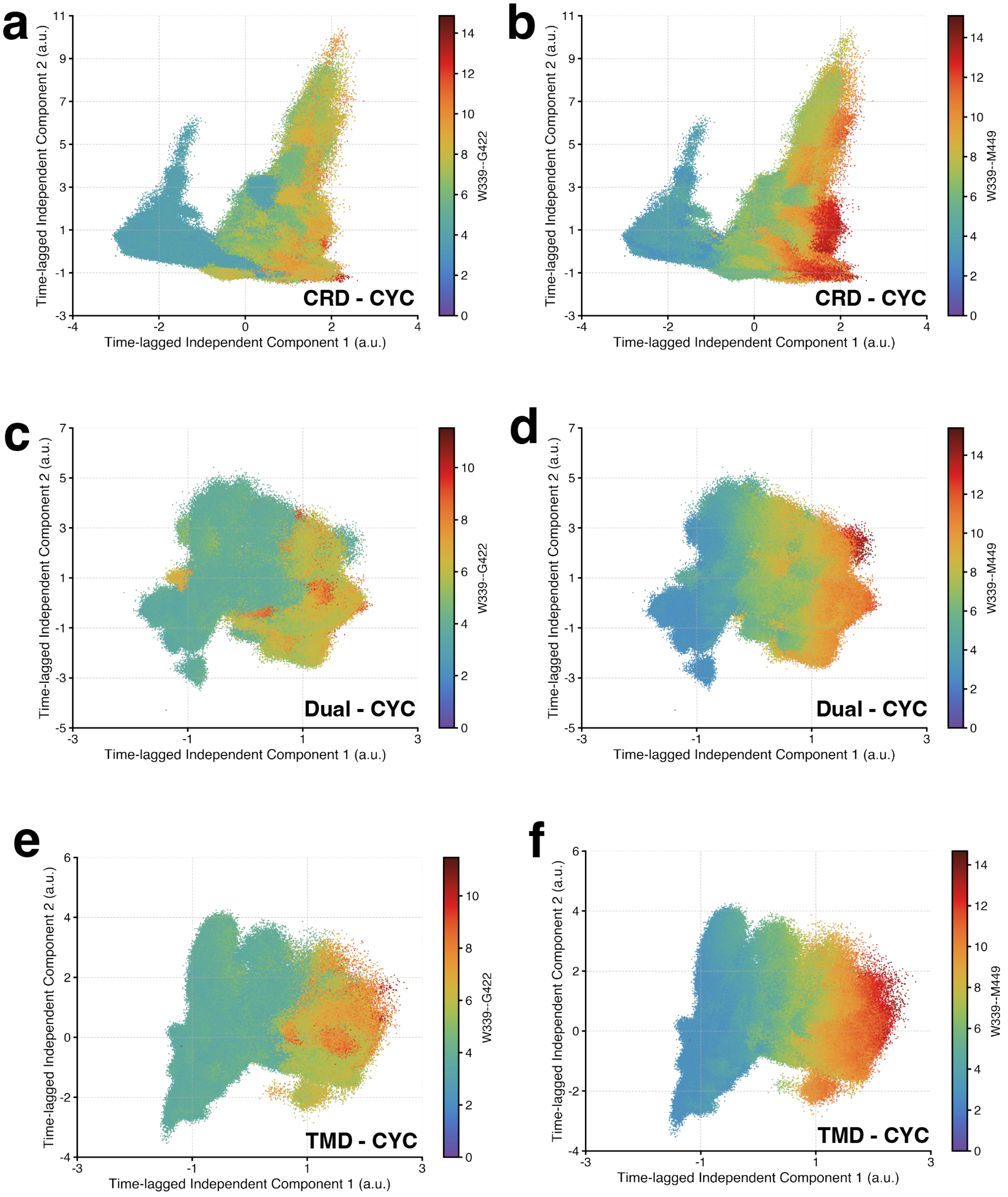


Figure S2: Projection of intracellular residue distances (W339$^{3.50f}$ - M449$^{6.30f}$, W339$^{3.50f}$ - G422$^{5.65f}$) onto the tICA space of (a,b) CRD-CYC, (c,d) Dual-CYC, and (e,f) TMD-CYC. In all cases, tIC1 is related with activation. This demonstrates that all systems can undergo activation through intracellular rearrangement of these residues.

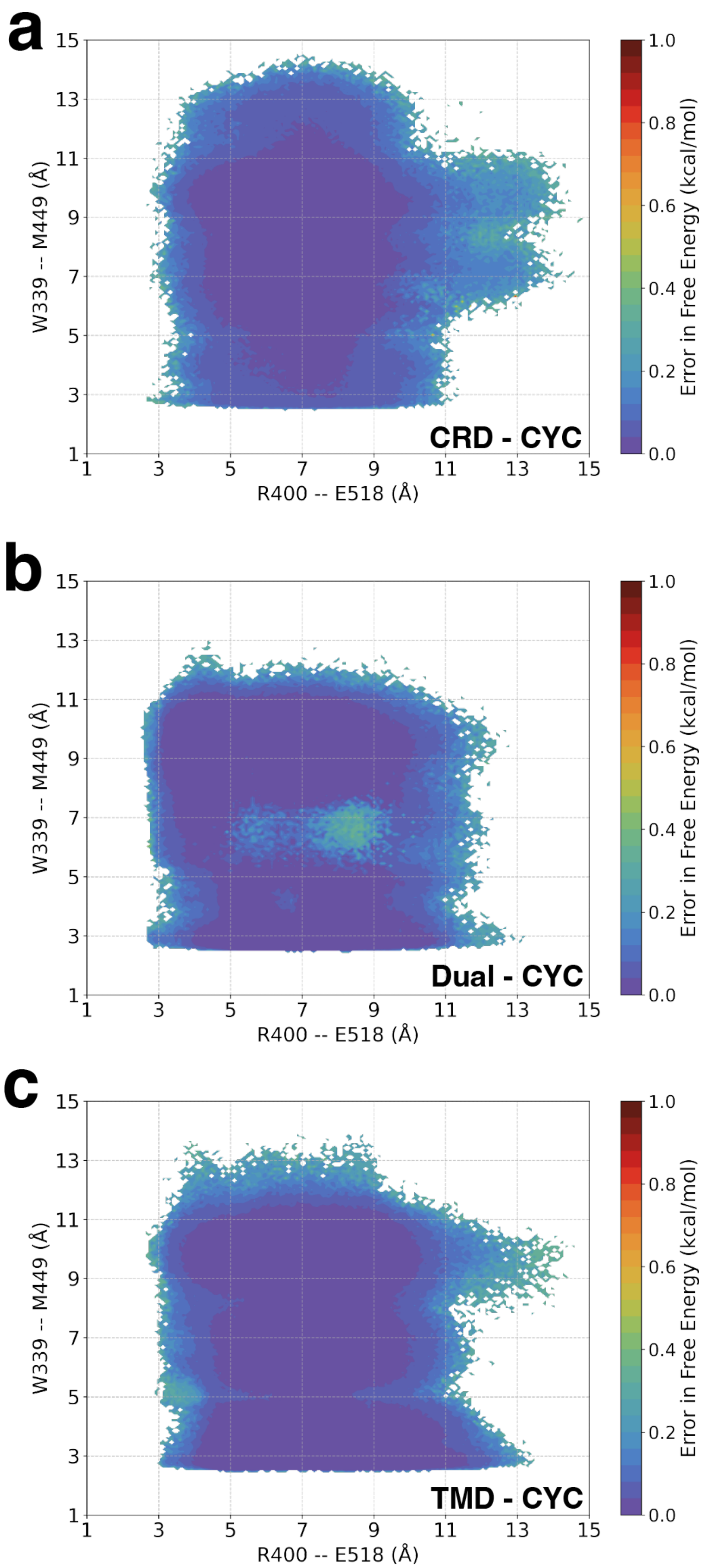


Figure S3: Errors in Figure 3 were calculated by using bootstrapping. 80% of the data was used in 200 iterations.

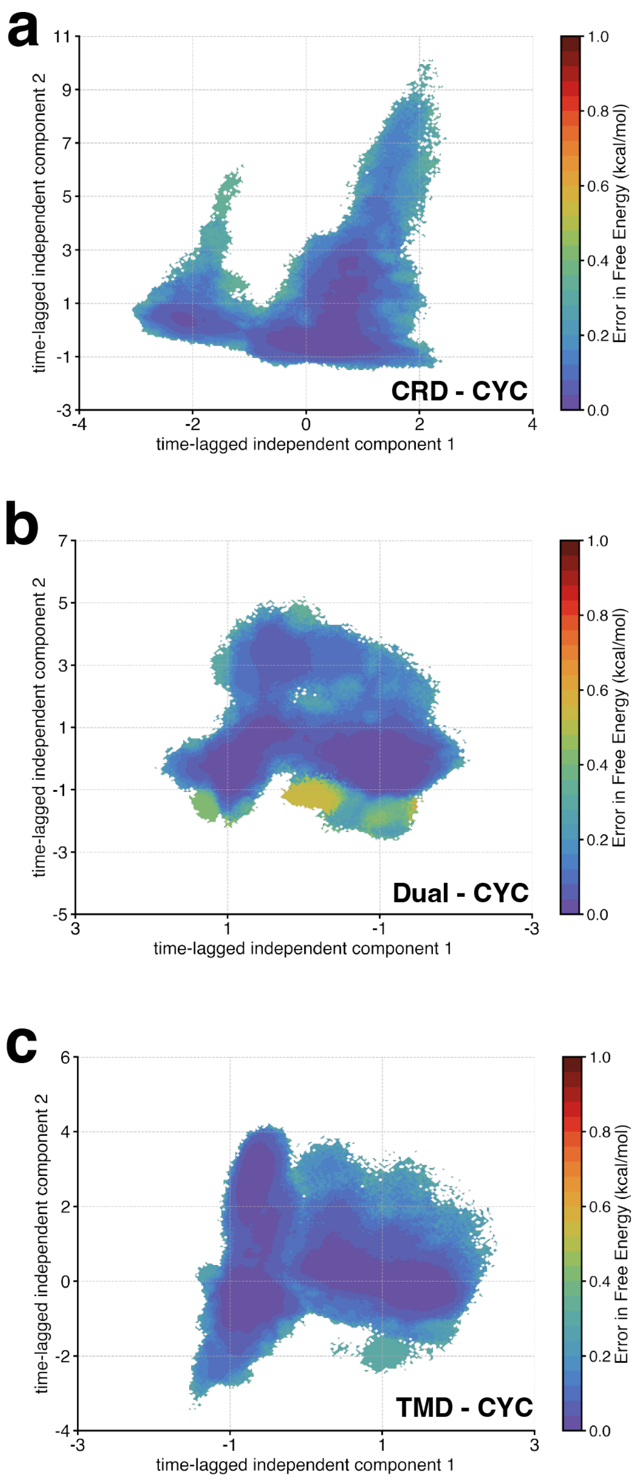


Figure S4: Errors in tICA plots in Figure 4 were calculated by using bootstrapping. 80% of the data was used in 200 iterations.

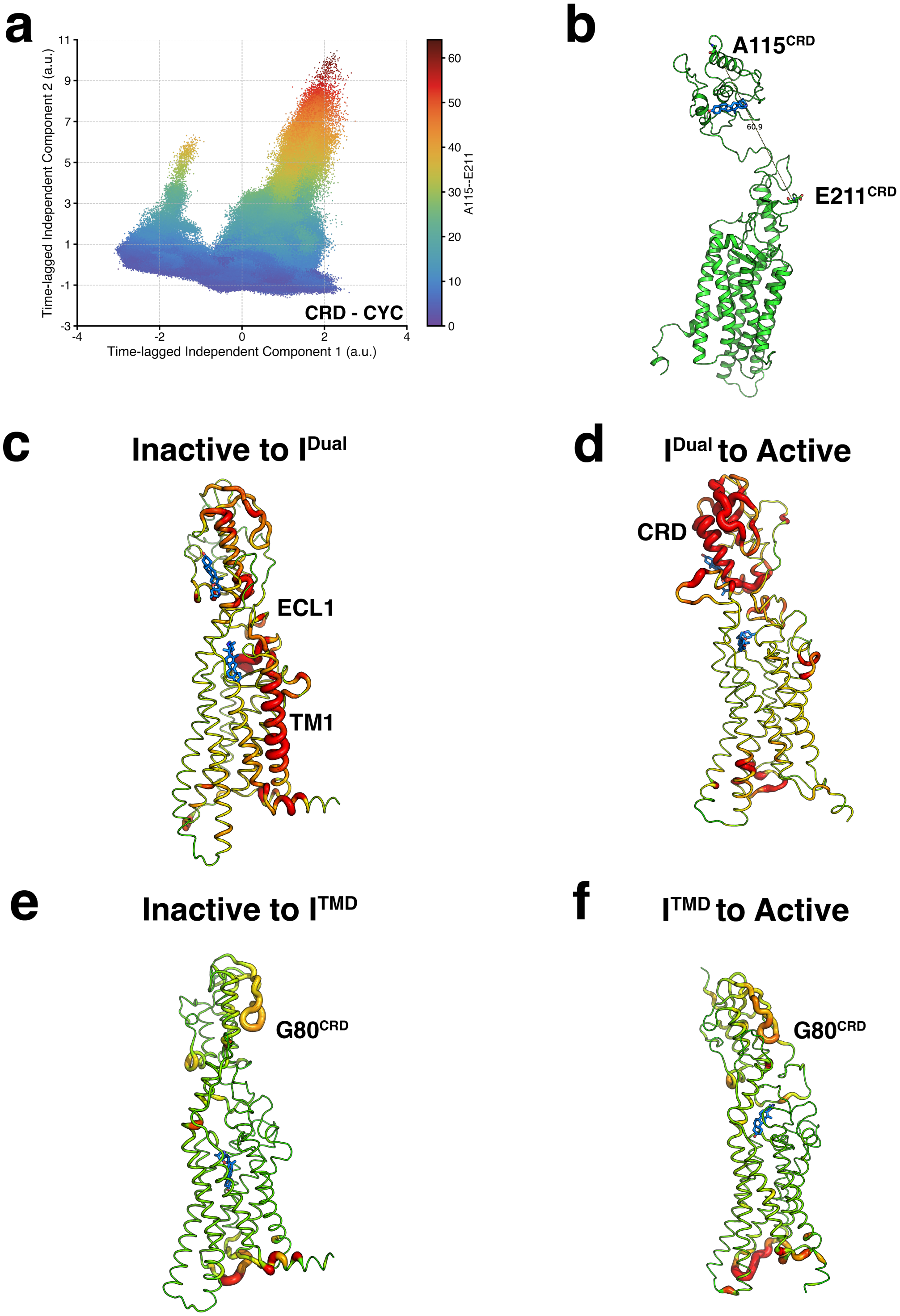


Figure S5: Projection of (a) A115$^{CRD}$ - E211$^{CRD}$ data onto the tICA space of CRD-CYC, (b) Obtained frame of CRD-CYC that has the highest tIC2 value of A115$^{CRD}$ - E211$^{CRD}$, (c) Transition state of Dual-CYC from inactive to I$^{Dual}$ - ECL1 and TM1 are major contributors. (d) Transition state of Dual-CYC from I$^{Dual}$ to active state - CRD is the major contributor. (e) Transition state of TMD-CYC from inactive to I$^{TMD}$ - G80$^{CRD}$ is the major contributor. (f) Transition state of TMD-CYC from I$^{TMD}$ to active state - G80$^{CRD}$ is the major contributor.

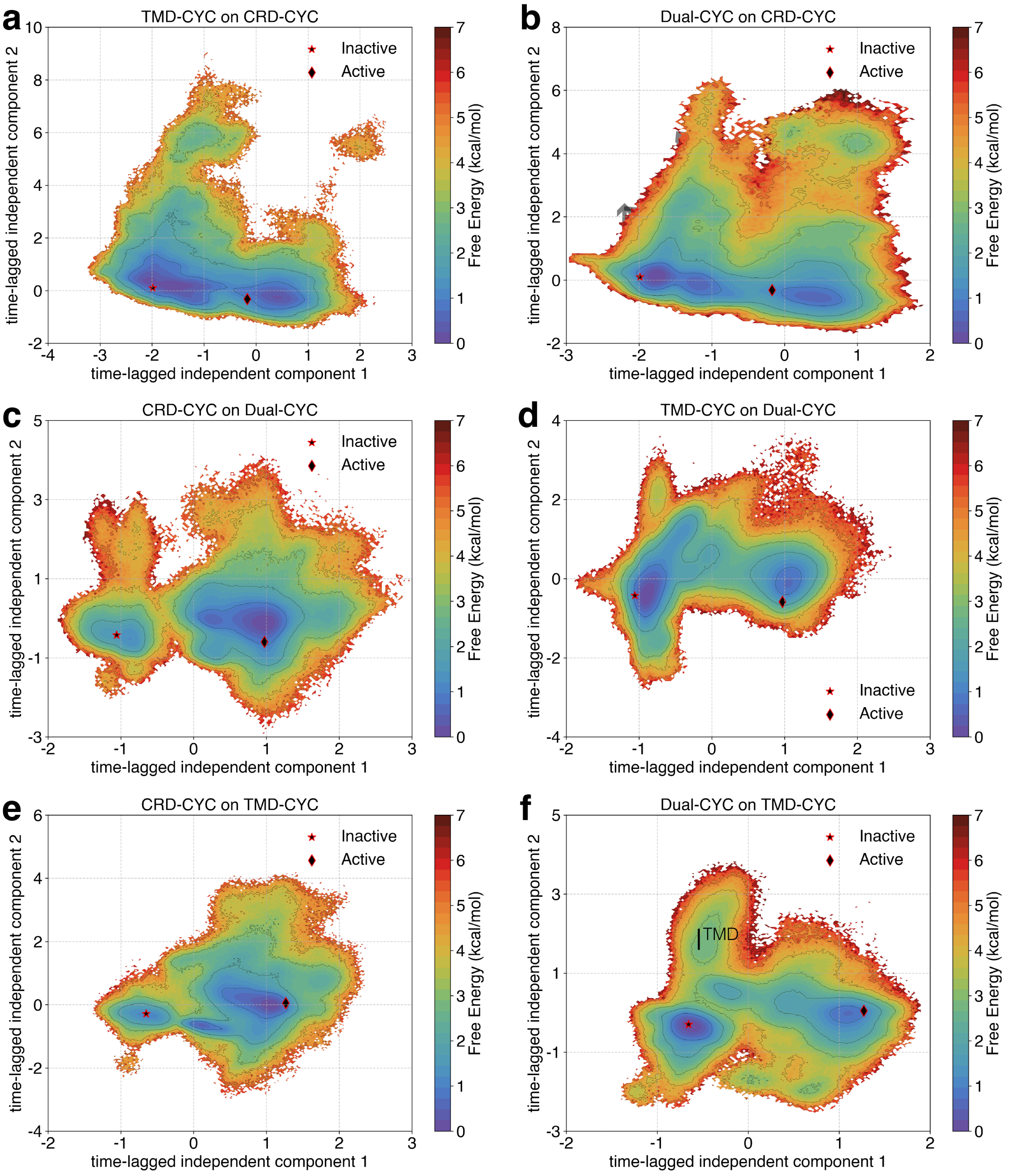


Figure S6: Projection of (a) TMD-CYC data onto the tICA space of CRD-CYC, (b) Dual-CYC data onto CRD-CYC, (c) CRD-CYC data onto Dual-CYC, (d) TMD-CYC data onto Dual-CYC, (e) CRD-CYC data onto TMD-CYC, (f) Dual-CYC data onto TMD-CYC. Figure (f) shows the presence of the intermediate state of TMD-CYC whenever TMD is occupied with cyclopamine. All plots are MSM weighted.

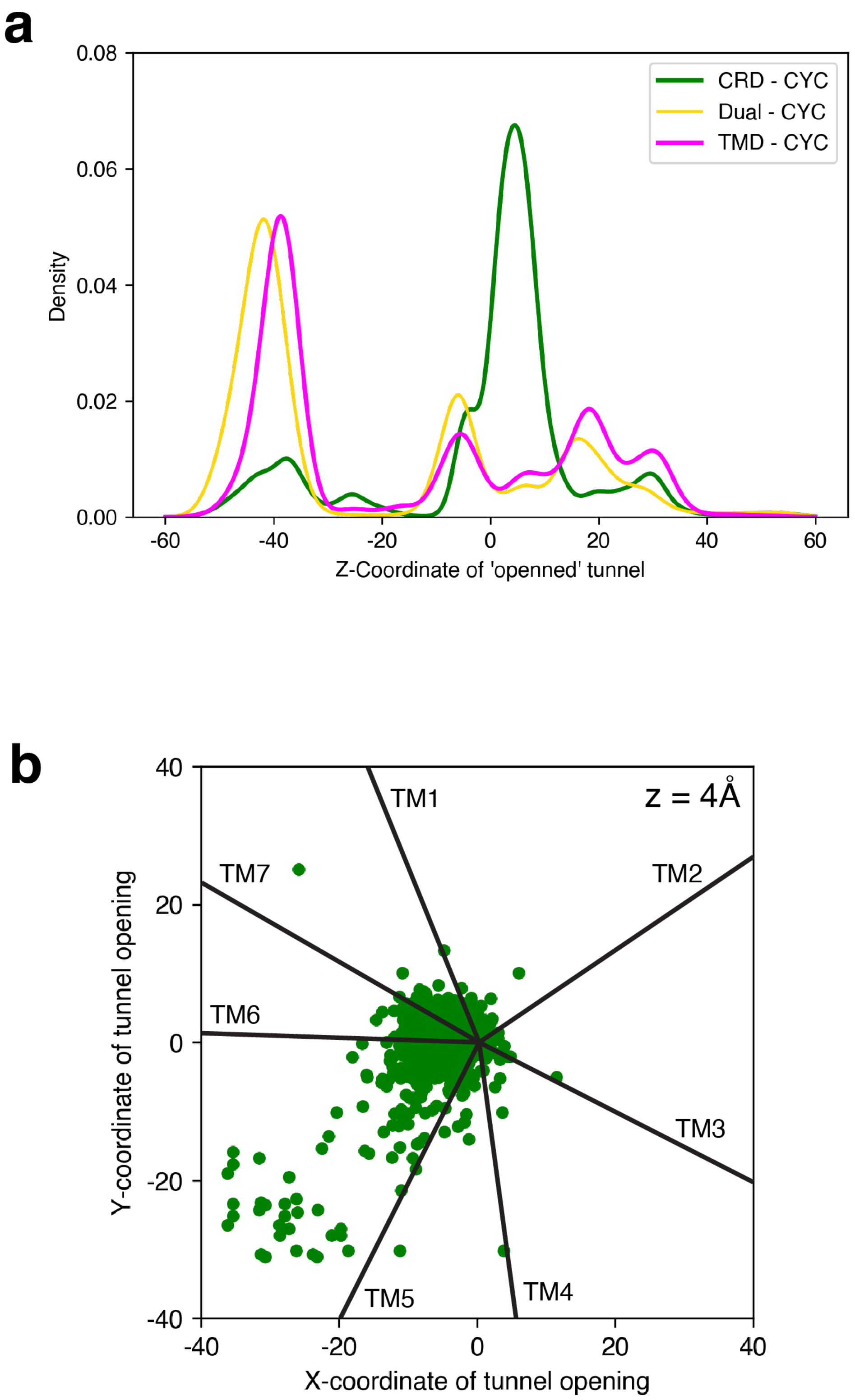


Figure S7: (a) Probability density of Z-coordinate of openned tunnel, where the diameter exceeds 5 angstroms. Simulations show that tunnel opens in the upper leaflet of the membrane at z = 4 Å. (b) The Y vs. X coordinate of the openned tunnel for CRD - CYC. Cluster is centered around the interface of TM5 and TM6. Transmembrane helical boundaries are indicated with black lines.

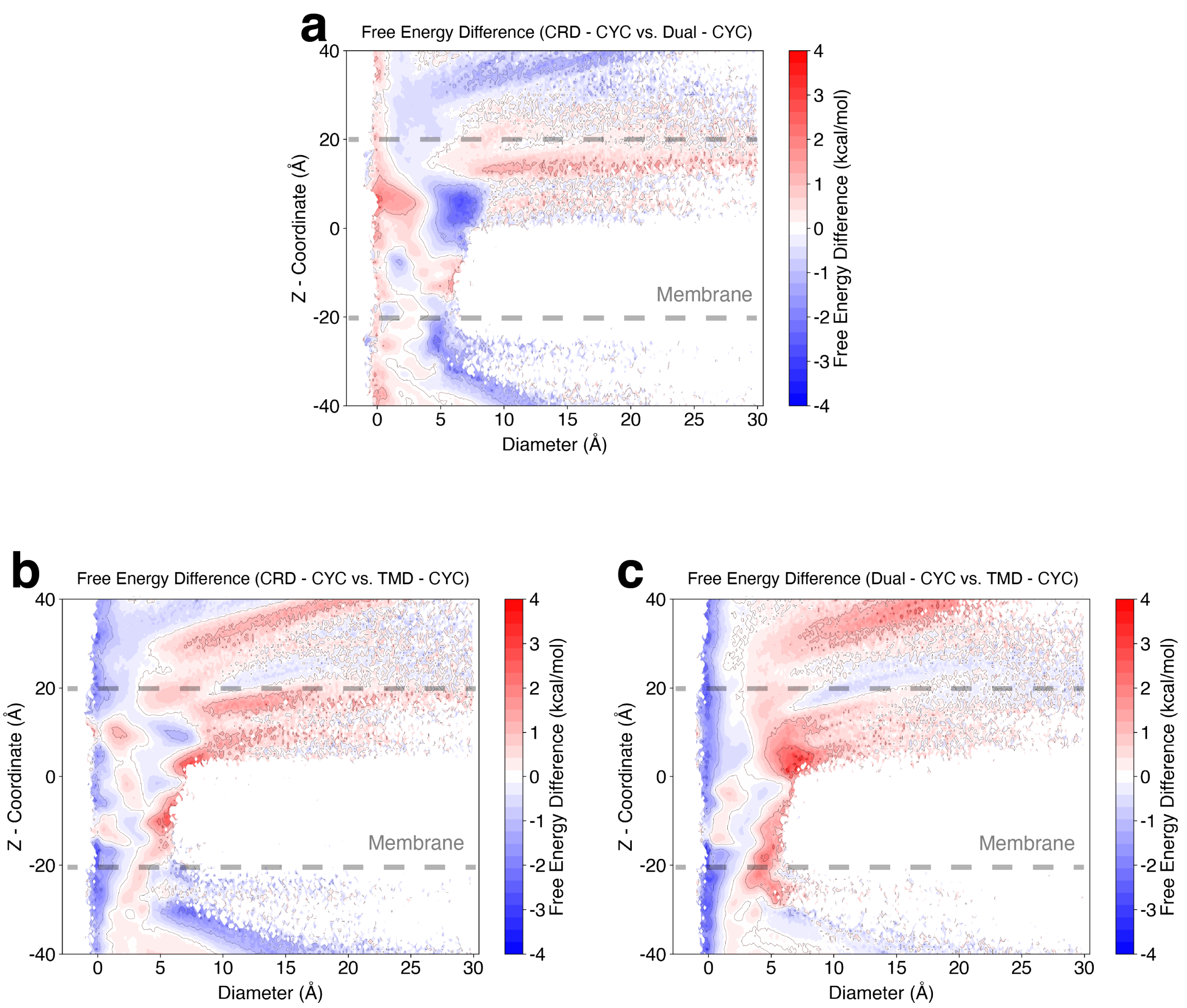


Figure S8: Free energy difference between the three plots shown in Fig 5. Free energy difference between (a) CRD - CYC and Dual - CYC, (b) CRD - CYC and TMD - CYC, (c) Dual - CYC and TMD - CYC. Red color clearly shows the expansion of tunnel for CRD - CYC in the upper leaflet.

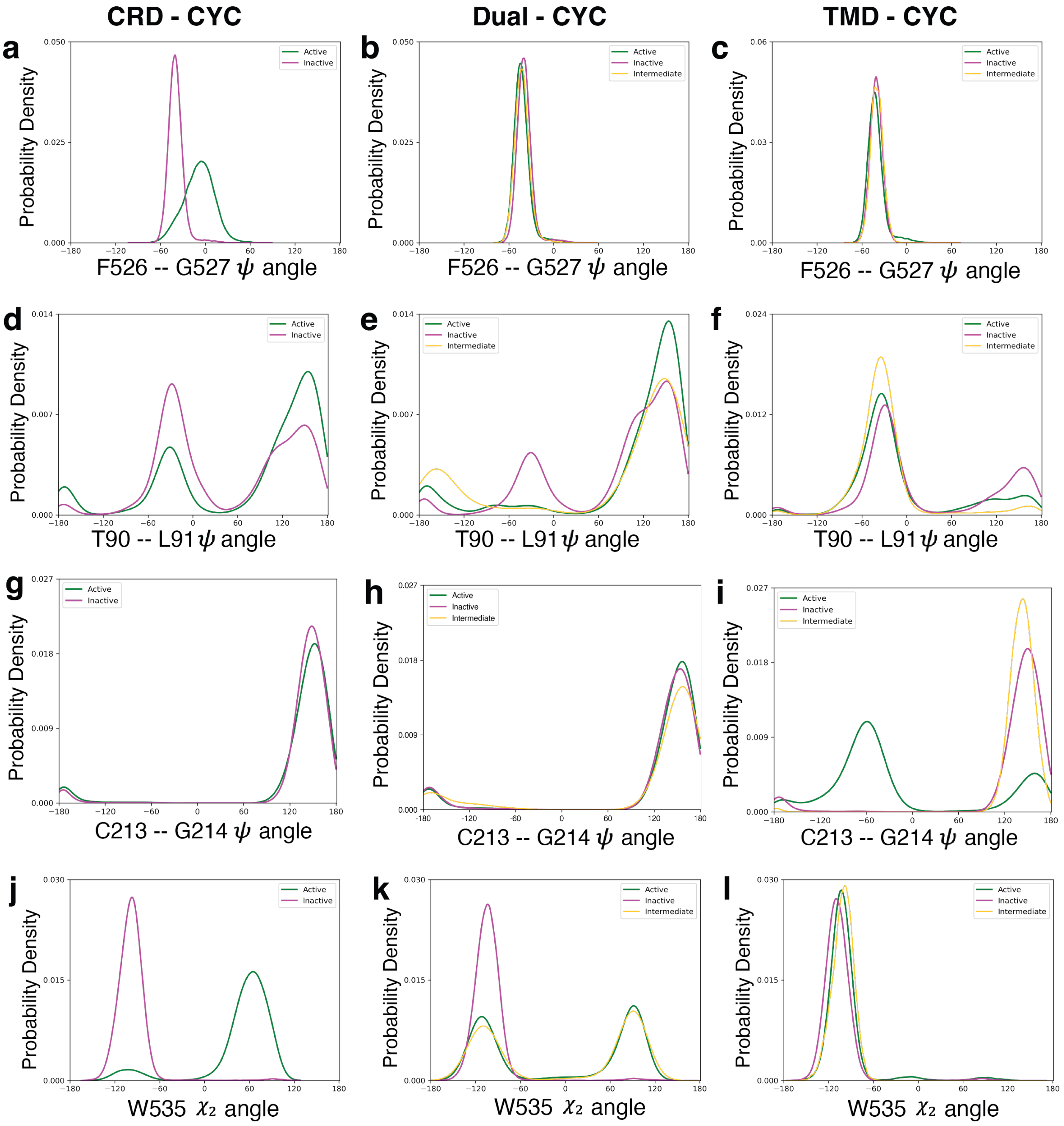


Figure S9: Probability density plots of $\psi$ dihedral angle rotation between F526$^{7.46f}$ and G527$^{7.47f}$ in (a) CRD-CYC, (b) Dual-CYC, (c) TMD-CYC. Probability density plots of $\psi$ dihedral angle rotation between T90$^{CRD}$ and L91$^{CRD}$ in (d) CRD-CYC, (e) Dual-CYC, (f) TMD-CYC. Probability density plots of $\psi$ dihedral angle rotation between C213$^{CRD}$ and G214$^{CRD}$ in (g) CRD-CYC, (h) Dual-CYC, (i) TMD-CYC. Probability density plots of $\chi_2$ dihedral angle rotation at W535$^{7.55f}$ in (j) CRD-CYC, (K) Dual-CYC, (l) TMD-CYC.

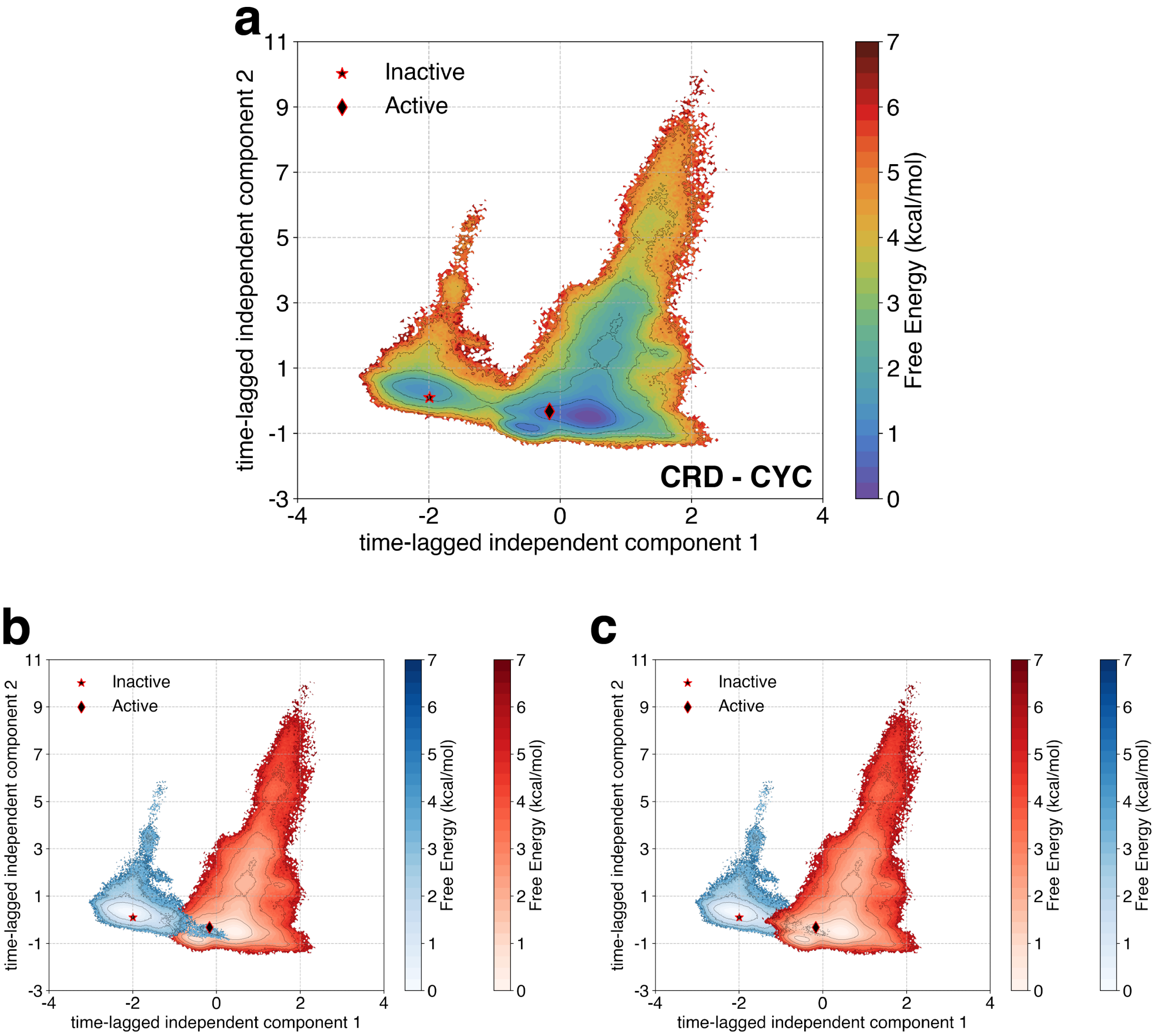


Figure S10: (a) Visualization of CRD - CYC projected onto the the tICA space, utilizing the two slowest components. The inactive and active starting points are marked. (b,c) The data for each inactive and active starting point are projected separately onto the tICA landscape. The overlap of two regions indicate the transitions from inactive to active state. The data from the inactive starting structure is depicted in blue, while the data from the active starting structure is shown in red.

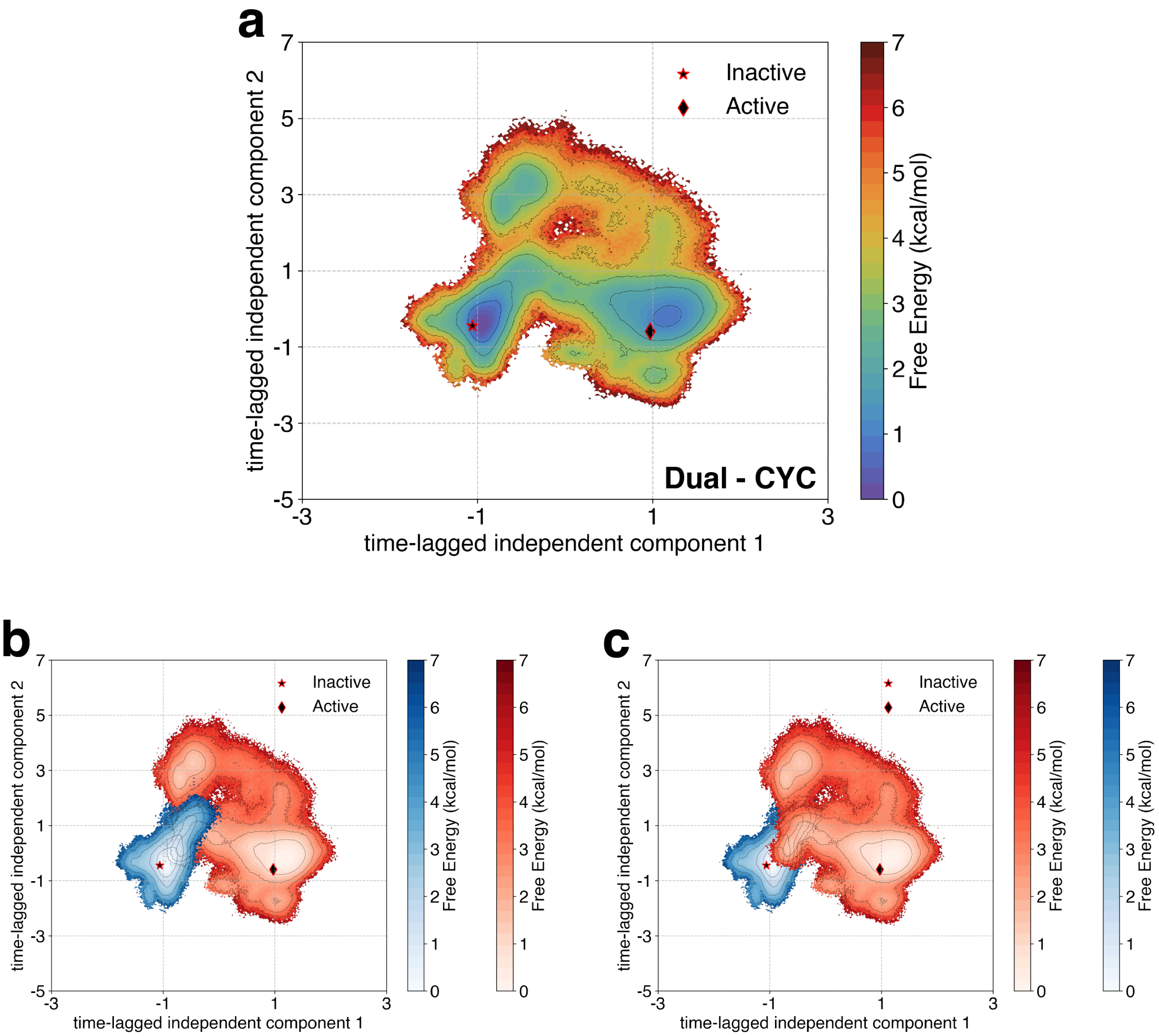


Figure S11: (a) Visualization of Dual - CYC onto the the tICA space, utilizing the two slowest components. The inactive and active starting points are marked. (b,c) The data for each inactive and active starting point are projected separately onto the tICA landscape. The overlap of two regions indicate the transitions from inactive to active state. The data from the inactive starting structure is depicted in blue, while the data from the active starting structure is shown in red.

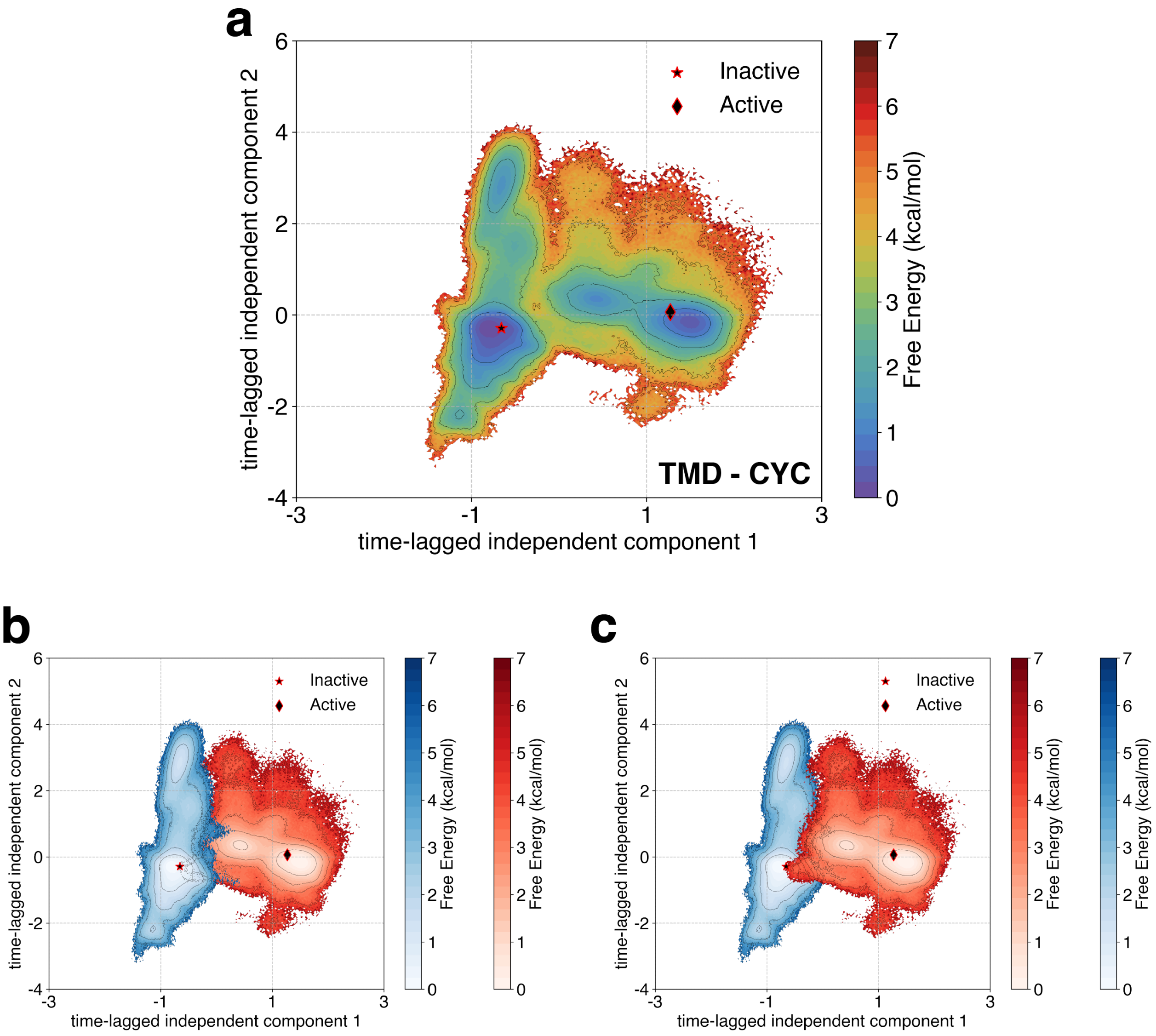


Figure S12: (a) Visualization of TMD - CYC projected onto the the tICA space, utilizing the two slowest components. The inactive and active starting points are marked. (b,c) The data for each inactive and active starting point are projected separately onto the tICA landscape. The overlap of two regions indicate the transitions from inactive to active state. The data from the inactive starting structure is depicted in blue, while the data from the active starting structure is shown in red.

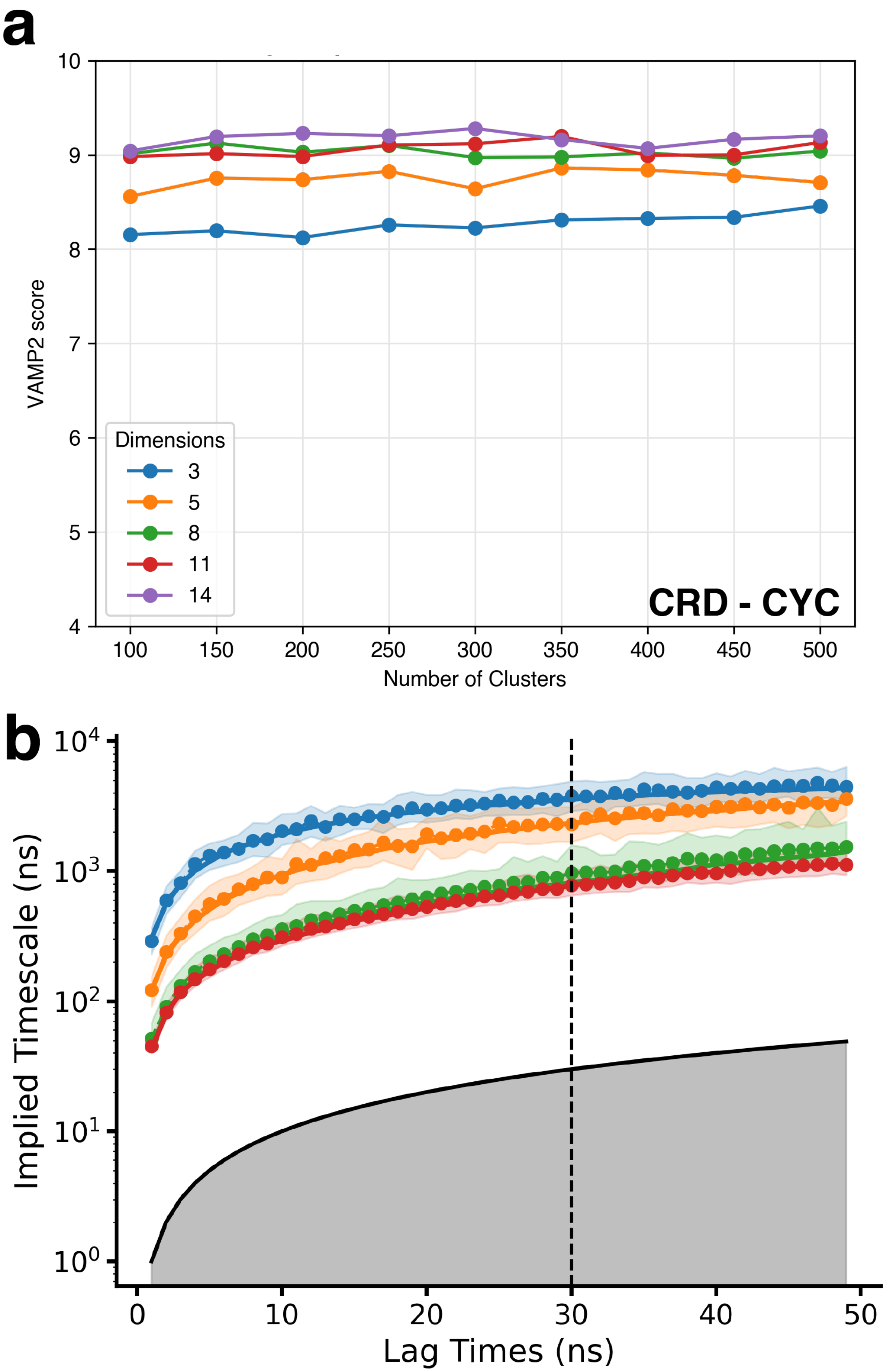


Figure S13: MSM construction for CRD - CYC. (a) VAMP2 score vs. Number of Clusters as a funtion of varying number of tICA components. (b) Implied Timescales vs. Lag Times for MSM with 150 clusters and 11 tICA components. The final MSM lagtime was chosen as 30 ns.

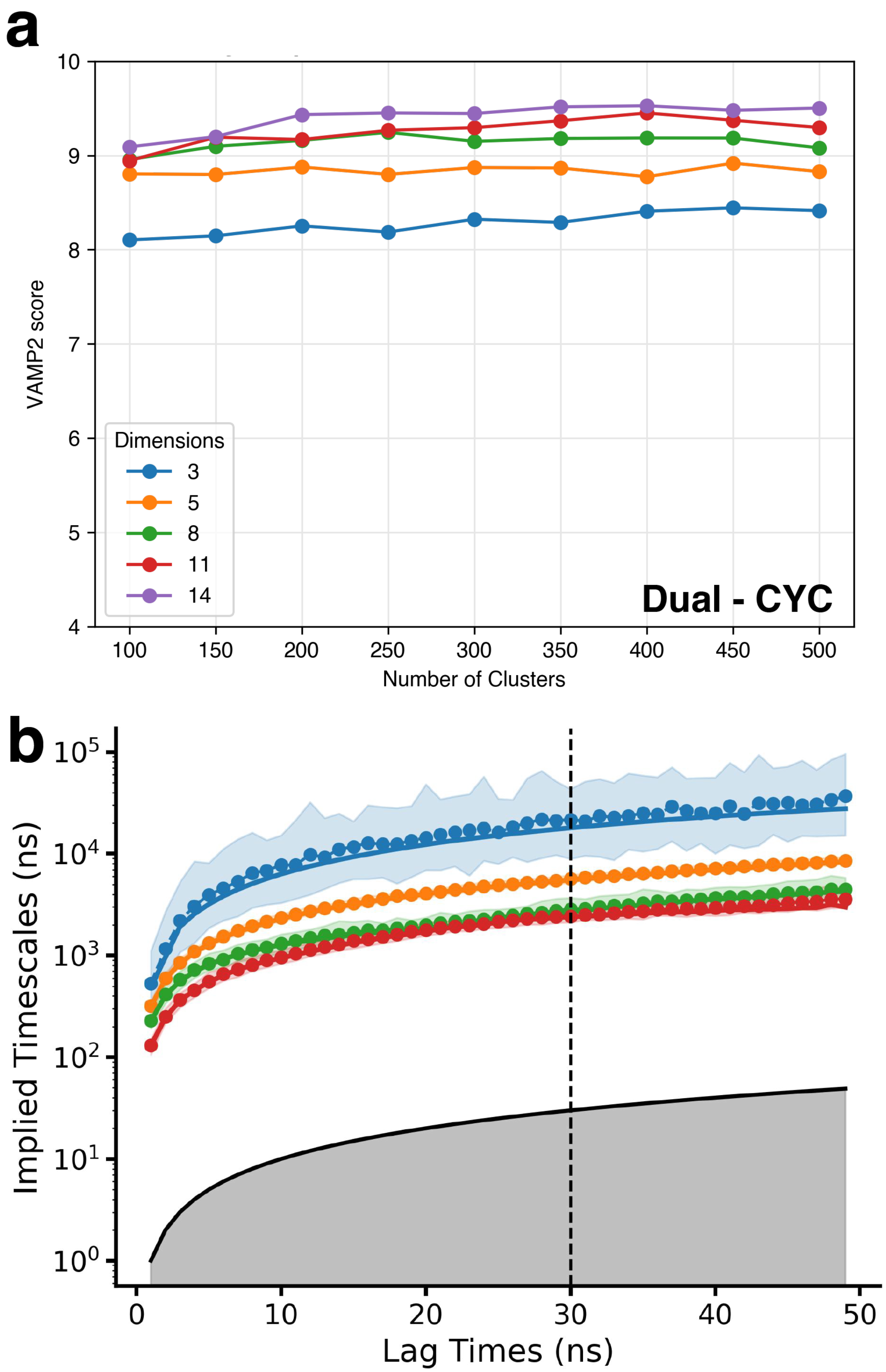


Figure S14: MSM construction for Dual - CYC. (a) VAMP2 score vs. Number of Clusters as a funtion of varying number of tICA components. (b) Implied Timescales vs. Lag Times for MSM with 400 clusters and 14 tICA components. The final MSM lagtime was chosen as 30 ns.

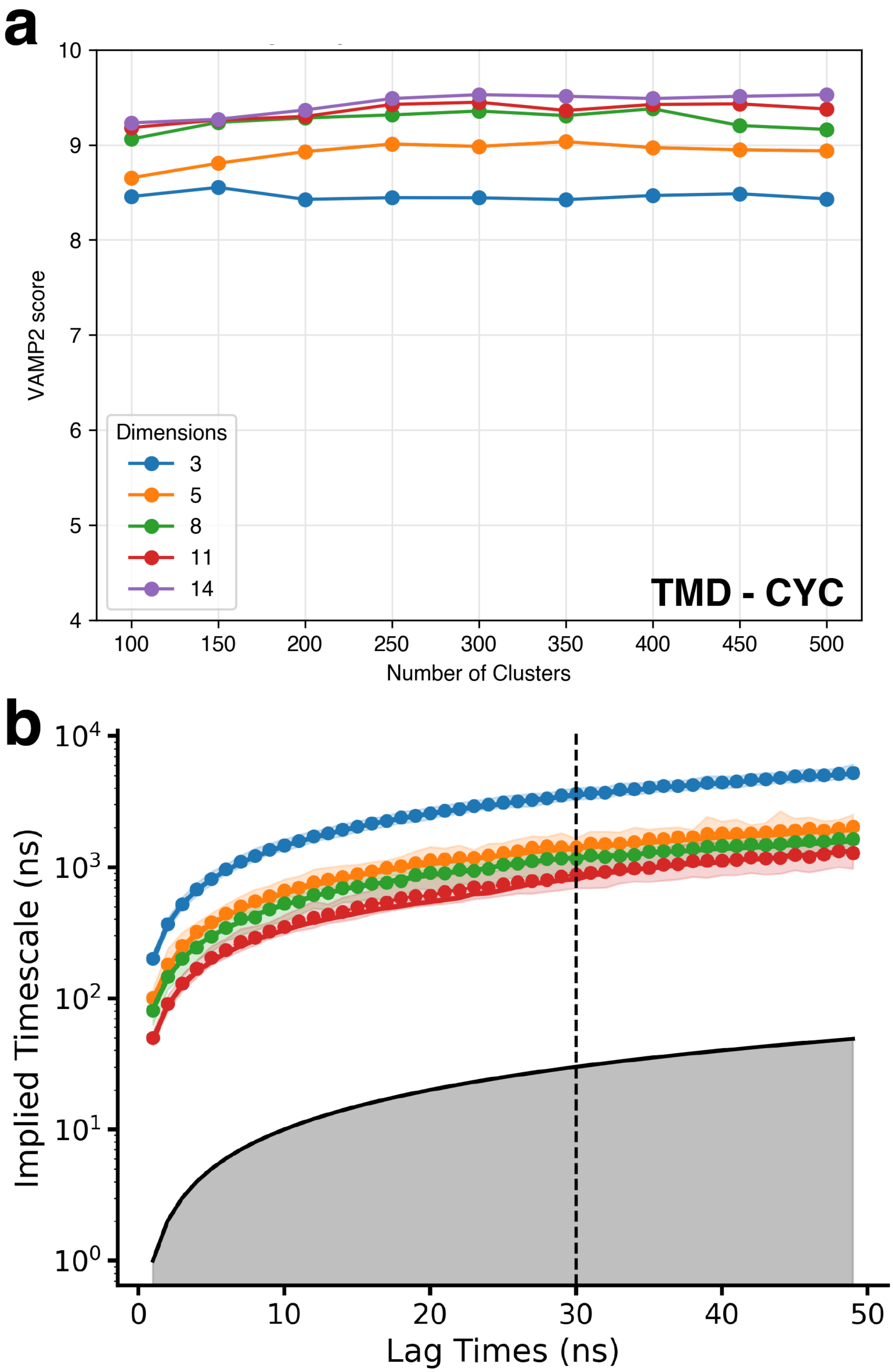


Figure S15: MSM construction for TMD - CYC. (a) VAMP2 score vs. Number of Clusters as a funtion of varying number of tICA components. (b) Implied Timescales vs. Lag Times for MSM with 150 clusters and 11 tICA components. The final MSM lagtime was chosen as 30 ns.

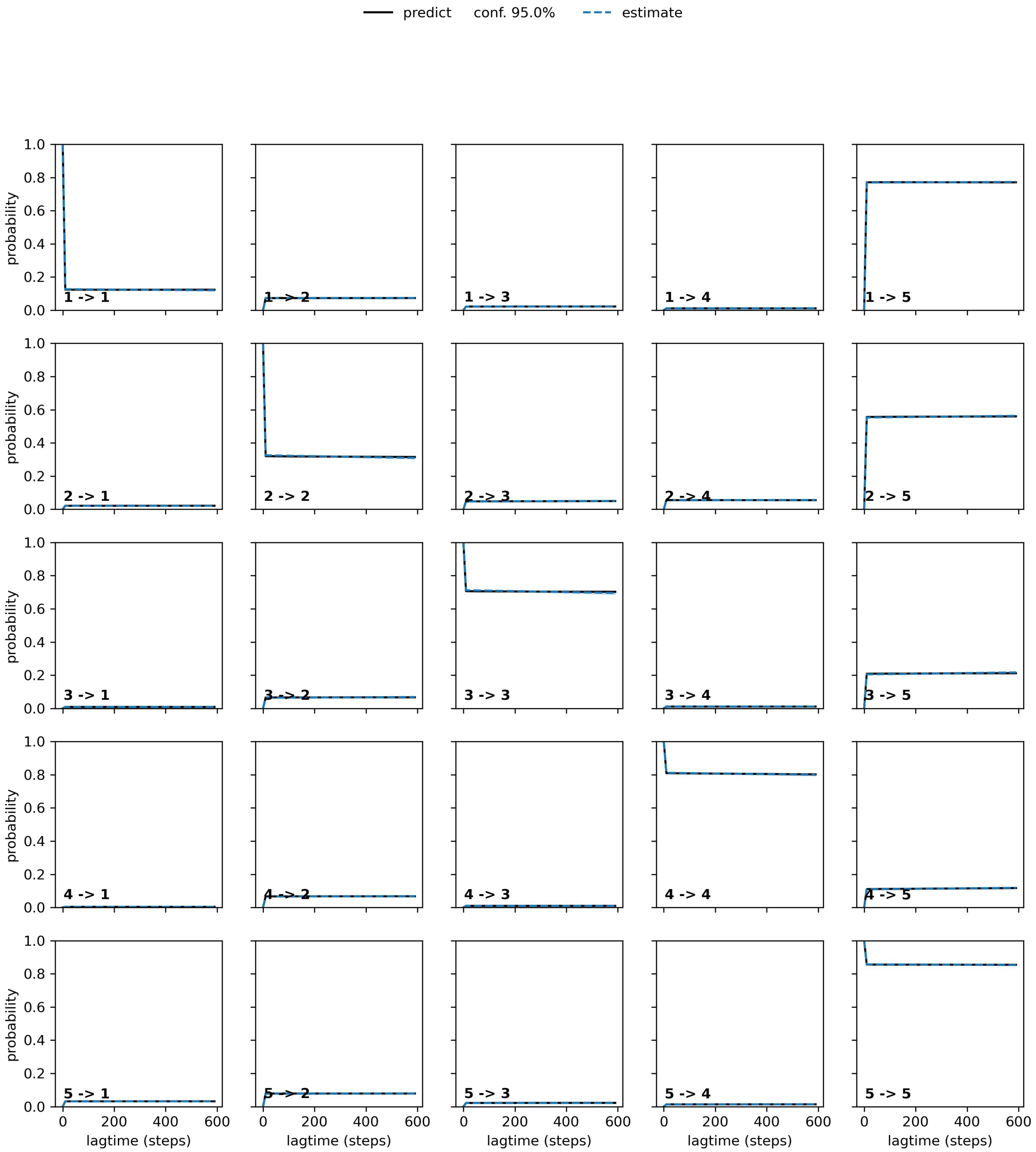


Figure S16: Chapman-Kolmogorov Test performed on CRD - CYC. The test is performed to validate the MSM with 150 clusters and 11 tICA components.

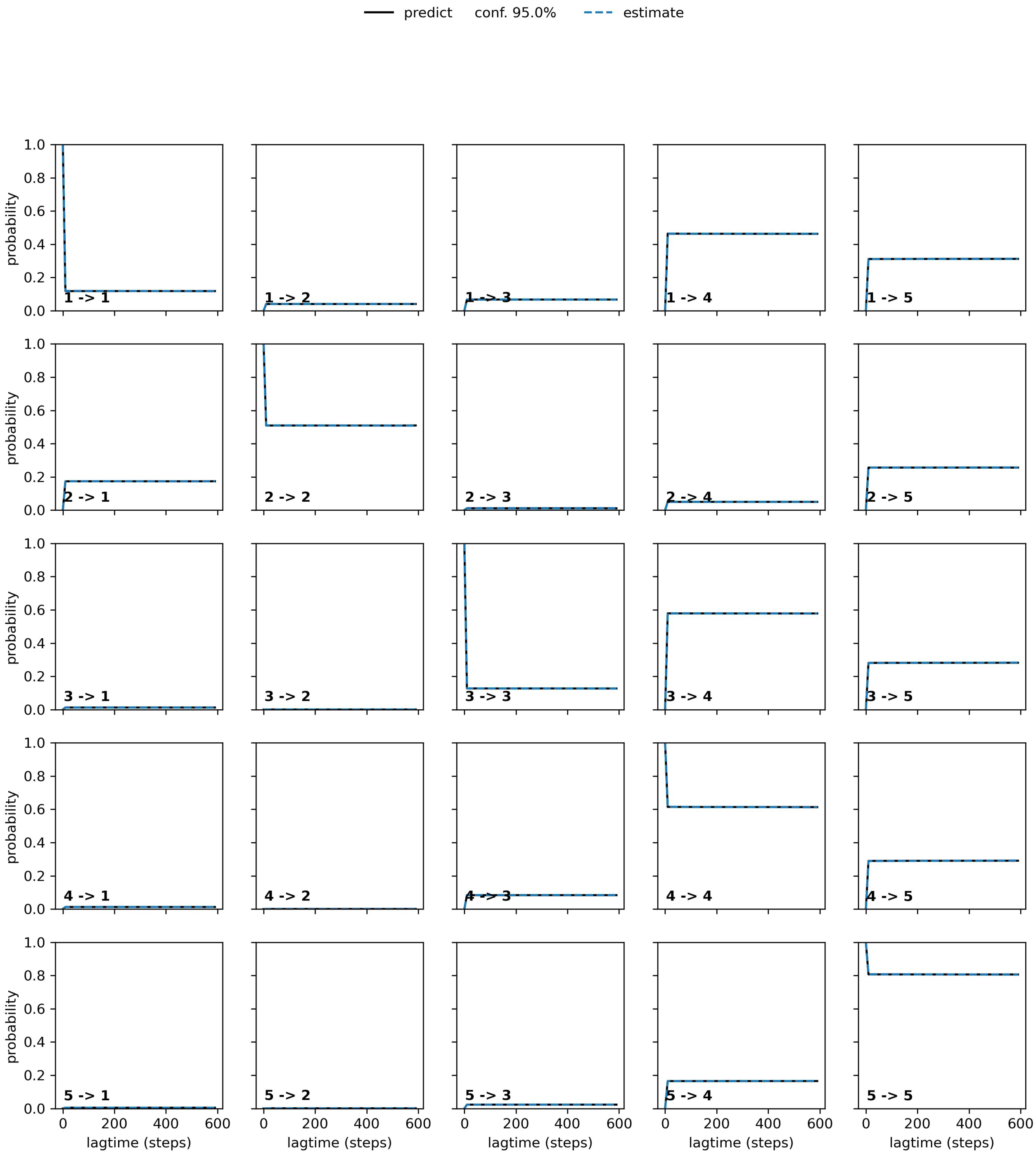


Figure S17: Chapman-Kolmogorov Test performed on Dual - CYC. The test is performed to validate the MSM with 400 clusters and 14 tICA components.

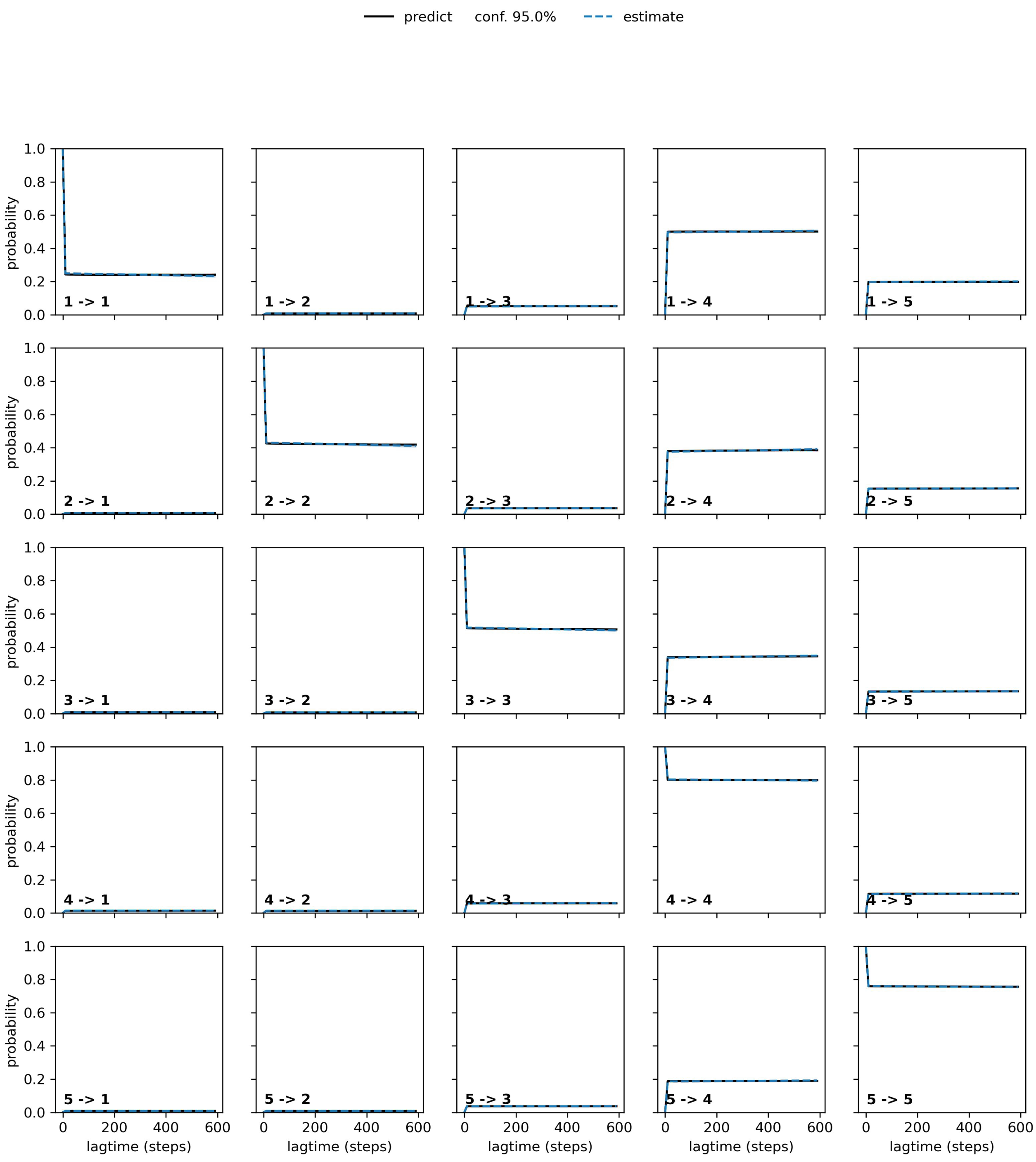


Figure S18: Chapman-Kolmogorov Test performed on TMD - CYC. The test is performed to validate the MSM with 150 clusters and 11 tICA components.

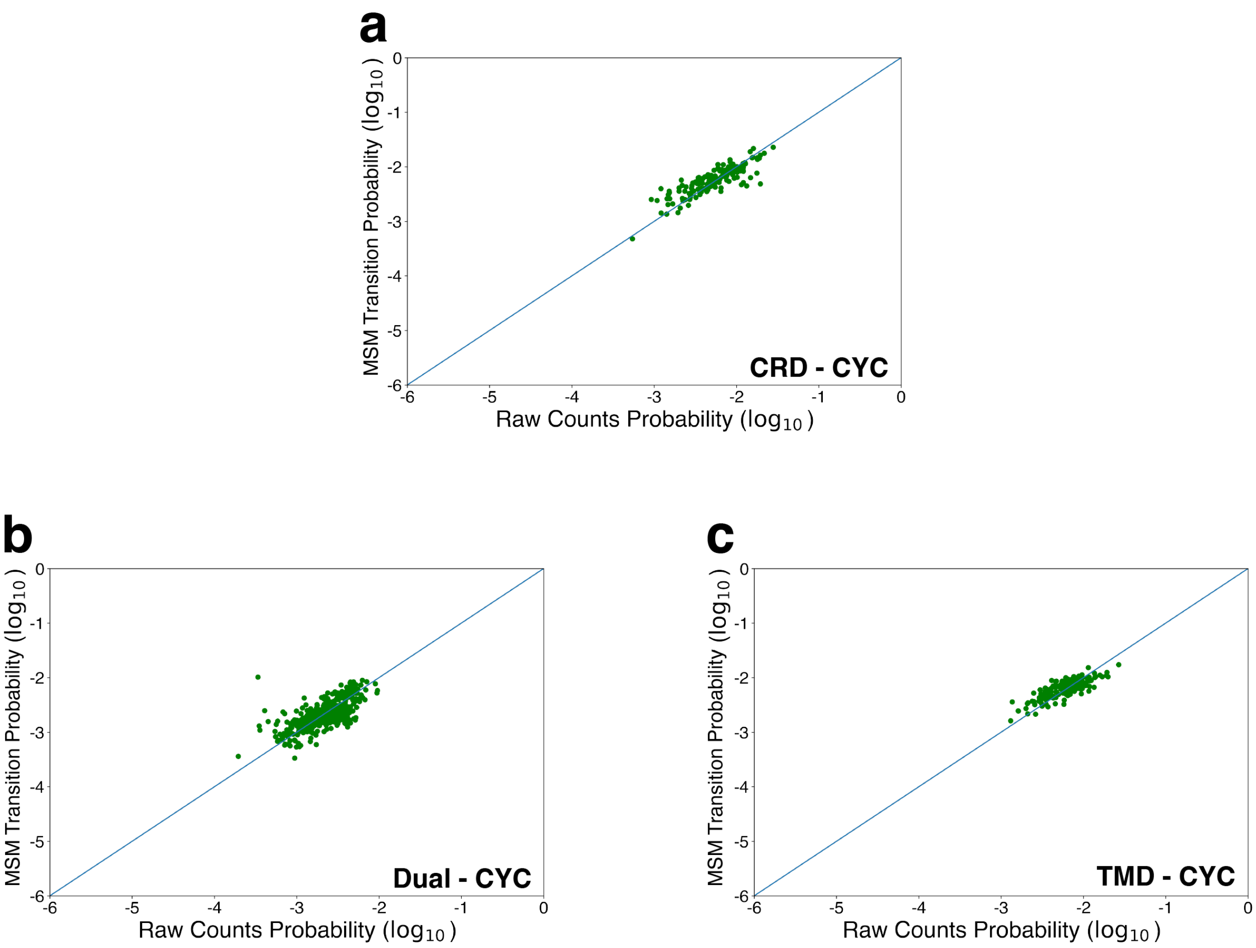


Figure S19: Raw counts versus MSM population for (a) CRD - CYC, (b) Dual - CYC, (c) TMD - CYC.

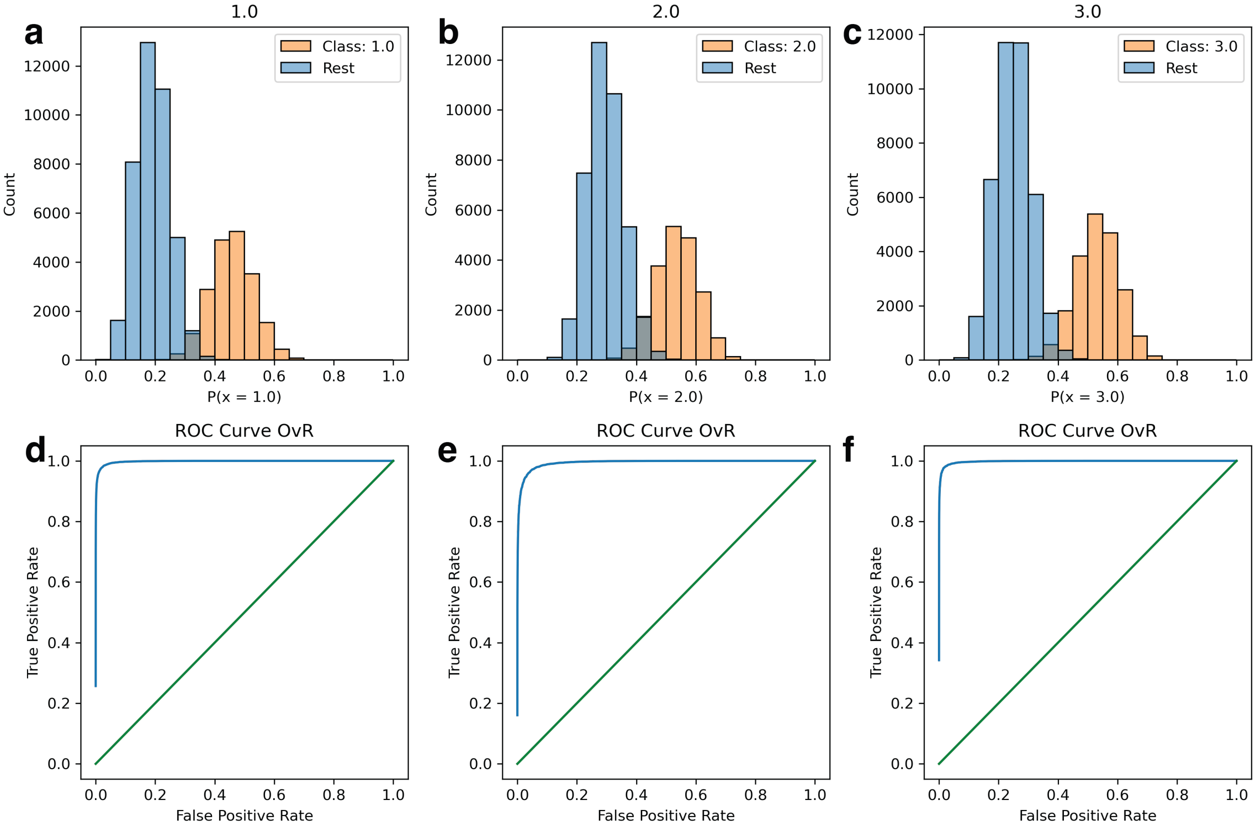


Figure S20: (a)-(c) Class separation Histograms for the 3 classes (a) CRD-CYC (b) Dual-CYC (c) TMD-CYC. The histograms were computed using a one-vs-rest (OvR) strategy. (d)-(f) Characteristic RoC Curves for the multiclass Random Forest Classifier. (d) CRD-CYC, (e) Dual-CYC (f) TMD-CYC. The curves show a classification accuracy greater than 99% for the supplied data.

Table S1: Modelled residues for inactive and active SMO.

| Modelled Residues | Constraints | Location |
|---|---|---|
| I429 | None | ICL3 |
| K430 | None | ICL3 |
| S431 | None | ICL3 |
| N432 | None | ICL3 |
| H433 | None | ICL3 |
| P434 | None | ICL3 |
| G435 | None | ICL3 |
| L436 | None | ICL3 |
| L437 | None | ICL3 |
| S438 | None | ICL3 |
| E439 | None | ICL3 |
| K440 | $\alpha$-helical | TM6 |
| A441 | $\alpha$-helical | TM6 |
| A442 | $\alpha$-helical | TM6 |
| S443 | $\alpha$-helical | TM6 |
| K444 | $\alpha$-helical | TM6 |
| I445 | $\alpha$-helical | TM6 |

Table S2: Membrane composition designed using CHARMM-GUI.

| Lipid | Upper Leaflet | Lower Leaflet |
|---|---|---|
| Cholesterol | 21 | 21 |
| POPC | 76 | 76 |
| Sphingomyelin | 4 | 4 |
| Total | 101 | 101 |

Table S3: 57 pairs of distances used for adaptive sampling. Double lines indicate separation of pairs of residues.

| | | | | | |
|---|---|---|---|---|---|
| P58 | P59 | R66 | R173 | Y85 | K133 |
| L106 | Y130 | L108 | W109 | W109 | L126 |
| R113 | W119 | A115 | E211 | R117 | W206 |
| R117 | G212 | W119 | Q123 | Q123 | F187 |
| R151 | W163 | W163 | R168 | C169 | F174 |
| N202 | W206 | W206 | E211 | W206 | G212 |
| Y207 | E208 | Y207 | D209 | F222 | L515 |
| H231 | F285 | H231 | R290 | L246 | F275 |
| F252 | S259 | F252 | F268 | W256 | F268 |
| Y262 | L353 | Q284 | R290 | G288 | E292 |
| R290 | R291 | F332 | A459 | L335 | W339 |
| W339 | G422 | W339 | M449 | W339 | G453 |
| W339 | W535 | F343 | M449 | L346 | I445 |
| L353 | F360 | K356 | F360 | Q380 | Y399 |
| V381 | V392 | Y397 | F474 | Y397 | Q477 |
| L419 | F457 | H433 | P434 | H433 | G435 |
| G435 | L436 | S443 | N446 | H470 | F474 |
| W480 | P513 | Y487 | Q491 | Y487 | I509 |
| Q502 | I504 | L516 | K519 | T534 | W537 |

Table S4: Round wise data collection for CRD - CYC.

| Simulation Round | Amount of Data for CRD - CYC |
|---|---|
| Round 1 | 200 |
| Round 2 | 100 |
| Round 3 | 100 |
| Round 4 | 100 |
| Round 5 | 100 |
| Round 6 | 100 |
| Total | 700 µs |

Table S5: Round wise data collection for Dual - CYC.

| Simulation Round | Amount of Data for Dual - CYC |
|---|---|
| Round 1 | 200 |
| Round 2 | 200 |
| Round 3 | 200 |
| Round 4 | 200 |
| Round 5 | 200 |
| Round 6 | 150 |
| Total | 1150 µs |

Table S6: Round wise data collection for TMD - CYC.

| Simulation Round | Amount of Data for TMD - CYC |
|---|---|
| Round 1 | 200 |
| Round 2 | 200 |
| Round 3 | 200 |
| Round 4 | 200 |
| Round 5 | 100 |
| Round 6 | 100 |
| Round 7 | 100 |
| Total | 1100 µs |
